# Antigen-specific Th17 T cells offset the age-related decline in durable T cell immunity

**DOI:** 10.1101/2025.06.05.658069

**Authors:** Ines Sturmlechner, Abhinav Jain, Jingjing Jiang, Hirohisa Okuyama, Yunmei Mu, Maryam Own, Cornelia M. Weyand, Jörg J. Goronzy

## Abstract

Older adults are susceptible to infections, in part due to waning of immune memory. To determine mechanisms that determine long-lasting versus short-term immunity, we examined varicella zoster virus (VZV) vaccination as a model system. We contrasted VZV antigen-specific T cells several years after vaccination in adults who had been vaccinated at young (<20 years) or older age (>50 years) with a live-attenuated vaccine that confers durable protection only when given at young age, or with an adjuvanted VZV component vaccine that elicits effective, long-lasting immunity in older adults. CD8^+^ T cells were highly sensitive to age-related changes showing T cell subset shifts, loss in TCR diversity and reduced stem-like features while gaining NK-like signatures without evidence for cellular senescence or exhaustion. VZV-specific CD4^+^ T cells were largely resilient to age and maintained phenotypic and TCR diversity. Immunization of older adults with the adjuvanted VZV vaccine did not reverse age-associated defects in CD8^+^ T cells. Instead, it selectively improved the functionality of VZV-specific Th17 CD4^+^ T cells and prevented their acquisition of Treg features, likely as consequence of lipid metabolic pathways. Collectively, our data indicate that effective vaccination in older adults is supported by the generation of a durable, antigen-specific CD4^+^ Th17 population that resists mis-differentiation into Tregs and that compensates for age-related defects in CD8^+^ T cells.

**One sentence summary:** Aging primarily impairs the CD8^+^ T cell VZV vaccine response, while effective VZV vaccination in older adults induces antigen-specific Th17 CD4^+^ T cells to compensate for aging defects.

## INTRODUCTION

Vaccination is one of the most successful interventions in modern medicine saving countless lives and improving public health globally. Childhood vaccinations have been transformative, however, infectious diseases remain a major cause of morbidity and mortality in older adults, when vaccinations are less efficacious. Numerous studies have tried to define the underlying age-related defects in the immune system, broadly referred to as immunosenescence(*1–3*). Successful vaccinations rely on the induction of immunologic memory that is mediated by B and T lymphocytes and plasma cells and typically manifest as the presence of antibodies in sufficient concentrations to neutralize the pathogen as well as the rapid T effector cell generation when the respective pathogen is encountered later in life(*4–6*).

Most previous studies have focused on memory generation during the first months after vaccinations by describing the increase in antibody titers and the frequencies of antigen-specific T follicular helper cells or cytokine-producing effector T cells as read-out systems(*7, 8*). Several immune defects in older age have been proposed, but many of the age-associated changes are relatively small and only incompletely explain the reduced clinical efficacy of vaccines in older adults. The responding T cell population is reduced in numbers and their ability to respond to T cell receptor (TCR) stimulation is blunted. DNA damage response pathways are increasingly compromised with aging(*9, 10*) which may exacerbate genomic stress and impair the rapid expansion of T cells on which T cell immunity highly relies on. While proliferating, T cells undergo fate decisions and differentiate into multiple phenotypic and functional T cell subsets. Fate decisions in older adults favor short-lived effector over longer-lived memory cells or follicular helper cells(*11*). Germinal center reactions involving the communication between B cells and follicular helper T (Tfh) cells are impaired, resulting in reduced generation of memory B cells and antibody-secreting, long-lived plasma cells(*12*). Fate trajectories are dynamic and develop over several months after vaccination(*13, 14*), a process that appears to be particularly vulnerable in the aging host with a failure to develop stem-like memory T cells.

A key determinant of successful vaccination is the durability of immune memory. Very few studies have examined the mechanisms determining long-term durability over years or decades even in young adults. Memory durability appears to differ depending on the vaccine type. A vaccine that induces remarkably durable immune memory is the live-attenuated yellow fever virus vaccine that confers life-long immunity(*15*). An example of differentially waning immunity was noted with the transition from the whole cell pertussis vaccine to the acellular vaccine(*16, 17*). The change in vaccination was associated with the resurgence of *Bordetella pertussis* infection among adolescents. The acellular vaccine induced mainly Th2 immune responses, while the whole cell vaccine induced mainly Th1 and Th17 responses that may account for superior and sustained protection. Similarly, the live-attenuated *Salmonella typhi* vaccines provide a longer duration of protection than the polysaccharide vaccine which may be due to a different balance between effector memory T cells and regulatory T cells(*18, 19*).

Here, we used varicella zoster virus (VZV) vaccination as a tool to uncover mechanisms of T cell memory durability depending on age and vaccine type. VZV typically causes chickenpox within the first few years of life while establishing latency. Since protection from VZV reactivation is mediated by T cells and not antibodies(*20*), we focused our analysis on T cells. Starting at about the age of 50 years, reactivation of the virus presenting as shingles is increasingly observed, culminating to 50% of older adults being affected by the age of 80 years(*21*). In 1995, the childhood VZV vaccine, Varivax, a live-attenuated virus was licensed in the US(*22*). Varivax requires a booster, but protects from chickenpox and, so far, prevents shingles(*23, 24*). The first vaccine used to protect older adults from shingles was Zostavax which employs the same virus strain as the childhood vaccination, however, at a ∼14-times higher viral dose. As opposed to the highly effective and durable response in children and young adults to the VZV vaccine, the effect of Zostavax is relatively short-lived with protection rates waning to ∼40% at 3-5 years post-vaccination(*25, 26*). In 2017, an adjuvanted VZV component vaccine was FDA-approved that is highly effective in preventing shingles even in >70 year-olds(*27*). Shingrix consists of recombinant VZV glycoprotein (gE) antigen and the AS01_B_ adjuvants and induces a highly protective immune memory in older adults with little waning within 10 years post-vaccination(*28*), a time window in which Zostavax protection has completely disappeared.

To pinpoint the molecular identity of T cell durability and efficacy, we leveraged the unique VZV vaccine landscape in the United States in 2021-2023 shortly after the introduction of Shingrix and discontinuation of Zostavax. This allowed us to recruit Shingrix and Zostavax vaccine recipients 3+ years after vaccination to directly compare the two contrasting vaccine T cell memories in older adults. In addition, we recruited younger adults having received Varivax, the same vaccine strain as Zostavax, in their childhood. We compared peripheral T cells specific for VZV gE, the antigen shared by all three vaccines. Via tri-modal single cell sequencing, we found that VZV gE-specific CD4^+^ and CD8^+^ T cells greatly differ in their susceptibility to age. Antigen-specific CD4^+^ T memory cells were largely stable in older Zostavax vaccinees except of a loss in the expression of interferon-related genes, however, without evidence for cellular senescence or exhaustion. In contrast, antigen-specific CD8^+^ T cells developed into end-differentiated, NK-like cells and contracted in TCR diversity, which may account for the loss in protective function. The more effective Shingrix vaccination could not restore these CD8^+^ T cell defects in older adults but selectively improved the functionality of VZV gE-specific CD4^+^ Th17 cells and prevented their degeneration into cells expressing Treg-related genes. Collectively, our data indicate that effective vaccination in older adults can be supported by the generation of a durable, antigen-specific CD4^+^ Th17 population that compensates for age-related defects in CD8^+^ T cells.

## RESULTS

### Influence of age on frequencies and heterogeneity of VZV-specific memory T cells

Vaccination with the live-attenuated VZV strain in children (Varivax) is protective for >20 years while the same vaccine strain in 50+ year-old adults (Zostavax) only induces partial and rapidly waning protection. To identify signatures in antigen-specific T cells that may account for the difference in protection and memory durability (Figure 1A), we recruited individuals who had received either the childhood (<20 years) or the adulthood vaccine (>55 years). Post-vaccination intervals were chosen for young adults (14 ± 1.7 years) and older adults (6 ± 0.8 years) after their last VZV vaccine dose. At the time of peripheral blood collection, these individuals were 24.3 ± 2.9 years (young adults, Y) and 75.5 ± 7.3 years old (older adults, O) (Supplementary Table 1).

**Figure 1:**
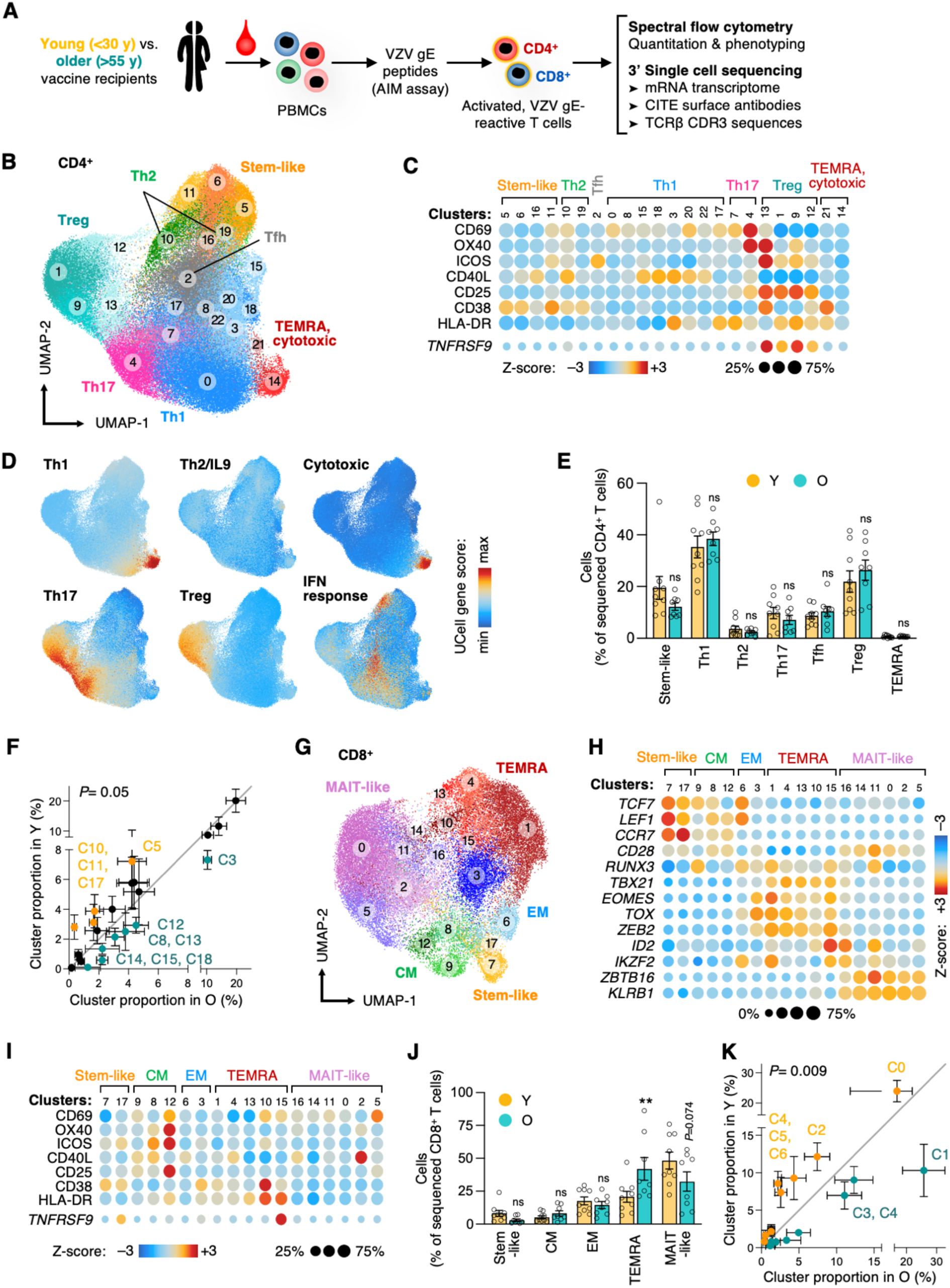
VZV glycoprotein E-specific CD8^+^ T cells are highly age-sensitive, while CD4^+^ memory T cells are resilient. **A**, Schematic of experimental design. **B,** Uniform manifold approximation and projection (UMAP) of VZV gE-reactive CD4^+^ T cells depicting distinct T cell clusters. **C**, Dotplot heatmap of activation markers via CITE protein (top) or gene expression (bottom) showing non-uniform activation marker expression across CD4^+^ T cell clusters. **D**, UCell gene score analysis of T helper signatures to annotate clusters to conventional T cell subsets. **E-F**, Relative subset (E) or cluster (F) frequencies of CD4^+^ VZV gE-reactive T cells comparing young and older vaccine recipients. Statistical analysis was performed by comparing probability vectors and permutation test. **G,** UMAP of VZV gE-reactive CD8^+^ T cells. **H,** Dotplot heatmap for VZV gE-responsive CD8+ T cells showing classical T cell subset markers. **I,** Dotplot heatmap of activation markers showing non-uniform activation marker utilization across CD8^+^ T cell clusters. **J-K,** Relative subset (J) or cluster (K) frequencies of CD8^+^ VZV gE-reactive T cells in Y and O vaccine recipients. Data show mean ± SEM (E,F,J,K). All datapoints represent distinct biological replicates. Data were compared by two-way ANOVA with Šídák’s multiple comparisons test (E,J). \*\**P*<0.01. ns, not significant.

We optimized activation-induced marker (AIM) assays(*29*) to comprehensively identify and collect VZV antigen-specific CD4^+^ and CD8^+^ T cells. Culture conditions were chosen to optimize expression of activation markers in peptide versus DMSO solvent control cultures and to identify the maximal number of antigen-specific T cells before cells divided ex vivo (Supplementary Figure 1A-E). For our AIM assays, we chose 42 hours after stimulation of peripheral blood mononuclear cells (PBMCs) with overlapping peptides of the VZV glycoprotein E (gE) as timepoint and CD69 and CD137 as the most inclusive activation markers for both CD4^+^ and CD8^+^ T cells as marker for FACS-mediated cell collection.

We found that net frequencies of VZV gE-reactive CD4^+^ or CD8^+^ T cells did not differ in older VZV vaccine recipients compared to younger vaccinees (Supplementary Figure 1F) despite the lower rate of shingles protection with age. To probe for qualitative changes in VZV gE-specific T cells, we subjected cells to single cell sequencing with 3 modalities: 3’ RNA and CITE surface antibodies, and *TRB* CDR3 sequences.

We profiled VZV gE-responsive CD4^+^ and CD8^+^ T cells from 9 young and 8 older live-attenuated VZV vaccine recipients by CITE-seq including 40 surface antibody features. We obtained 43,593 CD4^+^ and 19,350 CD8^+^ VZV gE-reactive T cells. CD4^+^ T cells were clustered into 23 subtypes as visualized by uniform manifold approximation and projection (UMAP, Figure 1B). Clustering resolution was chosen such that clusters subsetted well-defined conventional subsets. Almost all clusters included one to several activation markers in addition to CD69 (Figure 1C). Frequently, the expression of these activation markers was cluster-specific, validating our approach to use broad markers to capture cells for sequencing and use bioinformatic analyses to assess more specific combinations of activation markers. Subset annotation was performed using transcriptional signature gene scores (Figure 1D). Assessment of classical lineage-distinguishing cell surface molecules and transcription factors and transcripts confirmed the validity of the assignments and illustrating the cell heterogeneity of the VZV gE-specific CD4^+^ T cell response including subsets with features of stem-like, Th1, Th17, Treg, Th2, Tfh and TEMRA cells (Supplementary Figure 2A). The subsets Th1 (cluster, C0), Treg (C1) and Th17 (C4) were dominating in the VZV gE response which we confirmed by functional studies. To this end, we collected VZV gE-reactive CD4^+^ T cells by FACS based on the single cell sequencing-informed cell surface expression of CCR4, CCR6, IL7R, CD26 and TIGIT that distinguished these subsets (Supplementary Figure 2B). Purified cells were re-cultured, polyclonally stimulated with αCD3/αCD28 antibodies and profiled for cytokine secretion, cell expansion and survival (Supplementary Figure 2C-F). C1 (Treg) cells were predominantly FOXP3^+^ and IL7R^lo^, did not proliferate under these conditions, and were more prone to undergo apoptosis. In contrast, C0 (Th1) and C4 (Th17) cells showed high expansion potential, survival and differentially secreted high levels of cytokines including IFN-ψ, TNF⍺, and IL17A.

To determine whether the CD4^+^ T cell subset heterogeneity in the VZV gE response differs between young and older vaccine recipients, we compared the proportion of conventional subset and examined the distribution of antigen-specific CD4^+^ T cells in the high-dimensional clustering data and compared probability vectors of cluster frequencies by linear regression analyses. We found a trend towards significantly different subset distributions with age (*P*=0.05, Figure 1E,F) with several individual clusters marginally dominating in young (orange font such as C5) or older vaccine recipients (teal font such as C3). These data suggest that the VZV gE response of CD4^+^ T cells and their phenotypic heterogeneity is largely stable with age.

### CD8^+^ T cells specific for VZV gE are highly age-sensitive

Contrary to VZV gE-responsive CD4^+^ memory T cells, the antigen-specific CD8^+^ T cell response is subjected to extensive changes with age. Single cell clustering of VZV gE-responsive CD8^+^ T cells yielded 18 subsets (Figure 1G), differing in their profile of cell surface lineage molecules, the expression of activation markers, and the expression of lineage-determining transcription factors and targets (Figure 1H,I, Supplementary Figure 3A). The CD8^+^ response was dominated by TEMRA and MAIT-like cells, the latter showing a bias in TRAV-TRAJ usage characteristic of this subset (Supplementary Figure 3B). Young and older vaccinated adults differed substantially and significantly (*P*=0.009) in their preferred CD8^+^ cluster and cell type distributions (Figure 1J,K). Specifically, end-differentiated TEMRA subsets significantly increased with age, while stem-like and MAIT-like cells trended to decrease.

The sensitivity of CD8^+^ T cells to age was not restricted to phenotypic heterogeneity but also involved their TCR repertoire composition. While the VZV gE CD4^+^ T cell response was highly diverse with largely unchanged TCR diversity indexes and only a trend to increased clonality in older vaccine recipients, CD8^+^ T cells were again subject to considerable age-related defects (Figure 2A). TCRβ chain diversity in VZV gE-specific CD8^+^ T cells was nearly 10-fold lower than in CD4^+^ T cells even in young adults and further dramatically decreased in older vaccine recipients. Plotting the numbers of distinct VZV gE-reactive TCRβ chains ordered by decreasing clonal size versus the cumulative space they occupy illustrated the high degree of diversity of antigen-specific CD4 T cells (Figure 2B,C). Again, diversity was strongly contracted for CD8^+^ T cells but only mildly affected for CD4^+^ T cells of older individuals.

**Figure 2:**
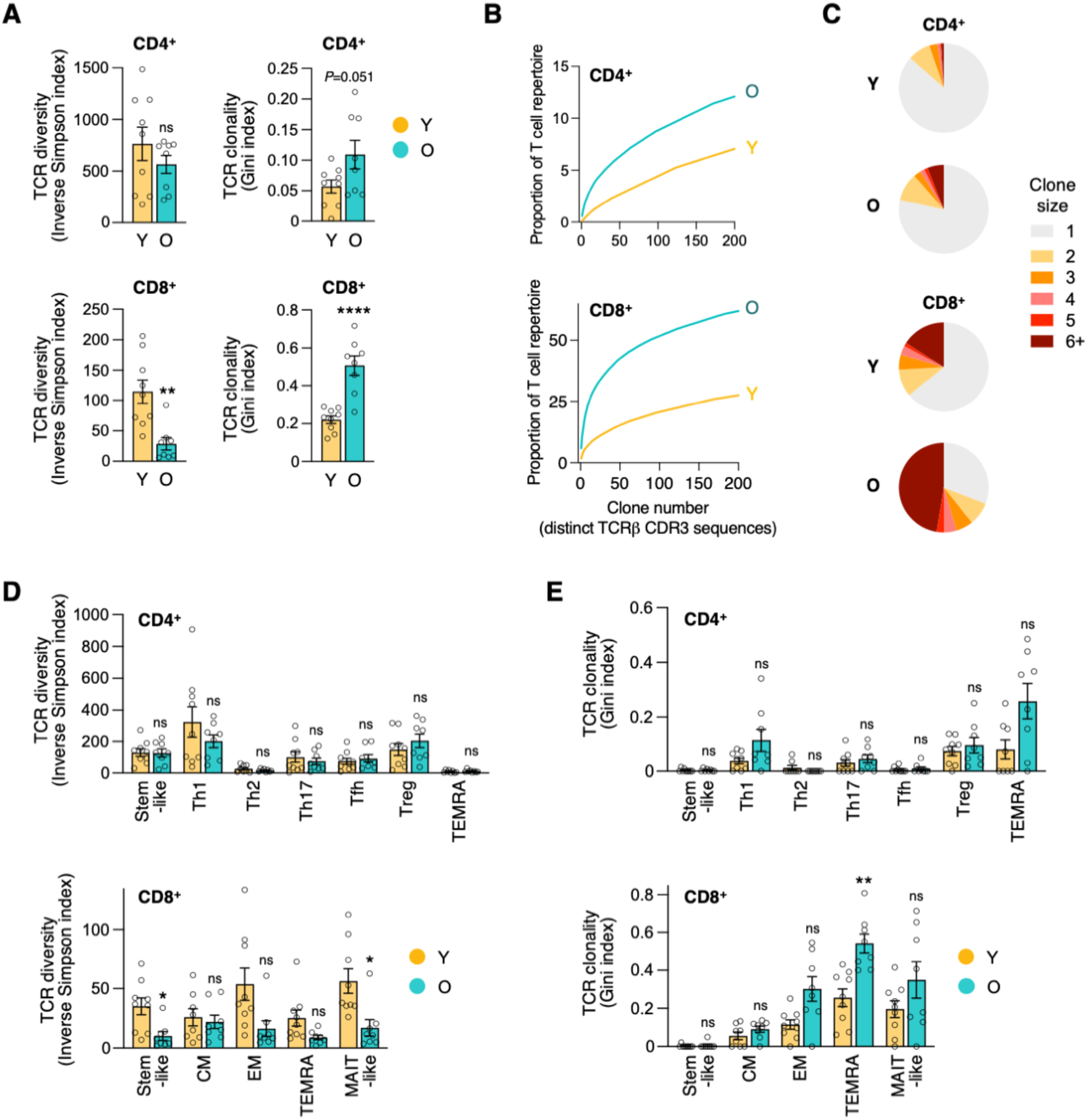
VZV gE-reactive CD4^+^ T cells maintain diversity with age while the repertoire of antigen-specific CD8^+^ T cells contracts. **A**, T cell receptor (TCR) diversity and clonality based on single cell TCRβ CDR3 sequences in VZV gE-reactive CD4^+^ and CD8^+^ T cells of young or older vaccine recipients. **B**, Cumulative TCR plots showing the TCRs ranked by the clone number in descending clone sizes vs the space they occupy. **C**, Pie chart of clonal expansion with size of the pie slices showing the proportion of clones of the indicated size. **D-E,** TCR diversity (D) and TCR clonality index (E) across T cell subsets of VZV gE-reactive CD4^+^ and CD8^+^ T cells in Y and O vaccine recipients. Data show the mean ± SEM (A,D,E). All datapoints represent distinct biological replicates. Data were compared by two-tailed, unpaired *t*-tests (A), or two-way ANOVA with Šídák’s multiple comparisons test (D,E). \**P*<0.05, \*\**P*<0.01, **** *P*<0.0001. ns, not significant.

The loss in CD8^+^ T cell diversity affected all subsets except the infrequent central memory cells, reaching significance for stem-like and MAIT-like T cells (Figure 2D). TCR clonality was increased in multiple subsets, predominantly in effector memory populations, and reached significance in the CD8^+^ TEMRA population of older adults (Figure 2E). These data indicate that reduced protection is unlikely to be attributed to differences in the frequencies and repertoires of antigen-specific CD4^+^ T cells, rather the age-associated contraction in CD8^+^ T cell diversity may increase the vulnerability of older adults.

### Increased IFN response in antigen-specific CD4^+^ T cells from young vaccine recipients

To probe for molecular signatures contributing to the age-related defect in T cell memory against VZV, we next assessed the gene expression profiles of memory T cell clusters. Dimension reduction of complex gene expression data using principal component analysis (PCA) showed a small shift in PC1 and PC2 of VZV gE-reactive CD4^+^ T cells from young and older vaccine recipients (Supplementary Data 4A). Pseudo-bulk differential expression analyses identified 579 up- and 535 downregulated genes in older adults. Subset clusters were unevenly affected with most differentially expressed genes (DEG) being found in Th1 (C0), Treg (C1) and stem-like (C5) cells (Figure 3A). Analysis of C1 Tregs did not identify enriched pathways. The stem-like C5 cluster lost self-renewing, stem-like-related transcription factors including *TCF4*, *LEF1* and *FOXO1* in older adults (Supplementary Figure 4B). Additionally, C5 DEGs in young adults showed an enrichment for type I and type II interferon responses, followed by increased NK cell-related features (Figure 3B). Conversely, C5 DEGs upregulated with age were enriched for pathways of translational and metabolic activity (Supplementary Figure 4C). An even larger enrichment for interferon responses at young age was seen for Th1 C0 and Th17 C4 cells (Figures 3B,C) suggesting that blunting of the interferon response with age is a common signature across multiple antigen-specific T cell subsets. This defect was not a sign of exhaustion of VZV gE- reactive CD4^+^ T cells as assessed by gene set enrichment analysis (GSEA) (Supplementary Figure 4D,E). Moreover, these T cells did not exhibit evidence of cellular senescence (Supplementary Figure 4F,G). Unexpectedly, CD4^+^ T cells of older vaccine recipients had gained innate lymphoid features, with their gene expression correlating with a gene set that had recently been described as designating the trajectory from CD4^+^ over CD8^+^, MAIT, iNKT, and γδ T cells to NK cells(*30*) (Figure 3D). These data suggest that, although largely stable in their numbers, TCR diversity and T cell subset phenotype, VZV gE-specific memory CD4^+^ T cells adapt their intrinsic response pathways with aging to shift towards NK-like, innate properties at the expense of potentially protective, anti-viral interferon signaling.

**Figure 3:**
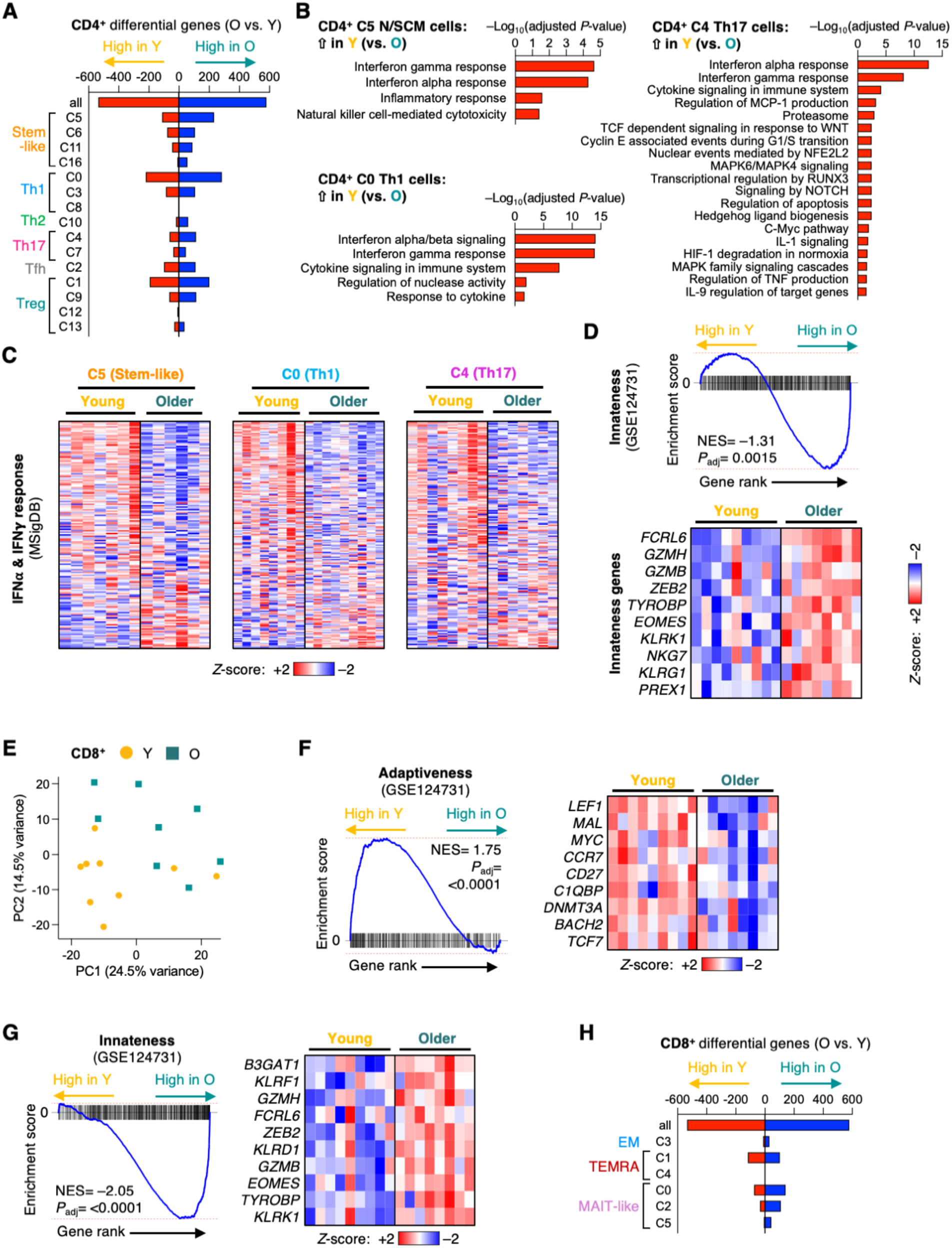
The type and extend of age-associated transcriptomic remodeling depends on the VZV gE-responsive T cell subset. **A**, Differentially expressed genes (DEG) in each CD4^+^ scRNA-seq cluster as identified by pseudo-bulk differential expression in O versus Y vaccine recipients. Only clusters with ≥2 male and ≥2 female subjects per group are included. **B**, Pathway enrichment for genes higher expressed in Y than in O in indicated CD4^+^ clusters. **C**, Gene expression heatmaps for gene sets “IFN⍺ response” plus “IFN𝛄 response” (MSigDB) based on pseudo-bulk expression comparing O versus Y vaccine recipients for 3 CD4^+^ T cell clusters (C5, C0, and C4). **D**, Gene set enrichment analysis (GSEA) comparing pseudobulk gene expression in CD4^+^ T cells from O versus Y vaccine recipients with “lymphoid innateness” gene set (GSE124731). Pseudo-bulk gene expression heatmap showing 10 genes from “innateness” gene cluster. NES, normalized enrichment score. **E**, Principal component analysis of CD8^+^ VZV gE-specific pseudo-bulk gene expression data for each vaccine recipient. **F-G**, GSEA and heatmaps for CD8^+^ VZV gE-specific T cells from O versus Y vaccine recipients. **H**, Differential gene expression in CD8^+^ VZV gE-specific T cells from O versus Y shown for total CD8^+^ T cells and indicated clusters.

### VZV gE-specific CD8^+^ T cells shift from adaptive to innate gene signatures with age

VZV gE-specific CD8^+^ T cells showed even more dramatic transcriptional changes with age, resulting in a clear separation of the gene expression profiles of young and older vaccine recipients by PCA (Figure 3E). GSEA identified a highly significant enrichment for adaptive immunity features in young and innate lymphoid immunity in older vaccine recipients (Figure 3F,G). In contrast, similar to the CD4^+^ T cell compartment, evidence for exhaustion and cellular senescence modules were low or absent (Supplementary Figure 5A-D). A total of 1,509 DEGs were identified in pseudobulk gene expression comparison of VZV gE-specific CD8^+^ T cells from young versus older vaccine recipients. These differences in gene expression were largely attributed to the shifts in subset distributions as described in Figure 1. Only a small number of DEGs were identified when pseudobulks for each cluster were compared (Figure 3H). Pathway analysis of these cluster-specific DEGs showed upregulation of translational processes in older MAIT-like cells and downregulation of signaling pathways in older TEMRA cells (Supplementary Figure 5E,F). Collectively, these analyses indicate that a shift of VZV gE-specific CD8^+^ T cells from self-renewing, stem-like T cell subsets to NK-like, innate subsets with age.

### VZV gE-specific Th17 response in older recipients of adjuvanted vaccines

An adjuvanted VZV gE component vaccine (Shingrix) that was licensed in 2017 in the US, provides efficient and long-term protection in older adults as opposed to the live-attenuated virus vaccine (Zostavax). To investigate how Shingrix elicits this exceptional protection despite age, we sought out to identify molecular determinants in T cells that may explain the higher durability of Shingrix. We recruited 13 individuals for single cell sequencing who had been vaccinated with Shingrix shortly after the introduction of the vaccine (ranging from 3 to 5 years ago). At the time of peripheral blood collection, Shingrix recipients were 69.5 ± 7.3 years old (referred to as Shingrix group, S), comparable to the previously described Zostavax recipients (as of now referred to as Zostavax group, Z) (Supplementary Tables 1, 2). We collected VZV gE-reactive T cells from Shingrix recipients by peptide stimulation of donor PBMCs and analyzed them by spectral flow cytometry and single cell sequencing (retaining 34,119 CD4^+^ and 9,989 CD8^+^ sequenced VZV gE-reactive T cells). In line with previous reports (*31*), the frequencies of VZV gE-specific CD4^+^ and CD8^+^ T cells were higher after Shingrix vaccination, irrespective of which activation marker was used (Figure 4A and Supplementary Figure 6A,B). In contrast, responses to an unrelated superantigen, SEB (*Staphylococcal* enterotoxin B), were not different between the two groups (Supplementary Figure 6C) suggesting similar immunological fitness of both vaccine recipient groups. Increased antigen-specific frequencies were not associated with a difference in TCR diversity or clonality for neither VZV gE-specific CD4^+^ nor CD8^+^ T cells, as concluded from Inverse Simpson and Gini indices (Figure 4B). Similarly, cumulative TCRβ-chain frequency plots showed no or minor differences with a slightly higher diversity and reduced clonality for VZV gE-specific CD4^+^ T cells of Shingrix recipients (Supplementary Figure 6D).

**Figure 4:**
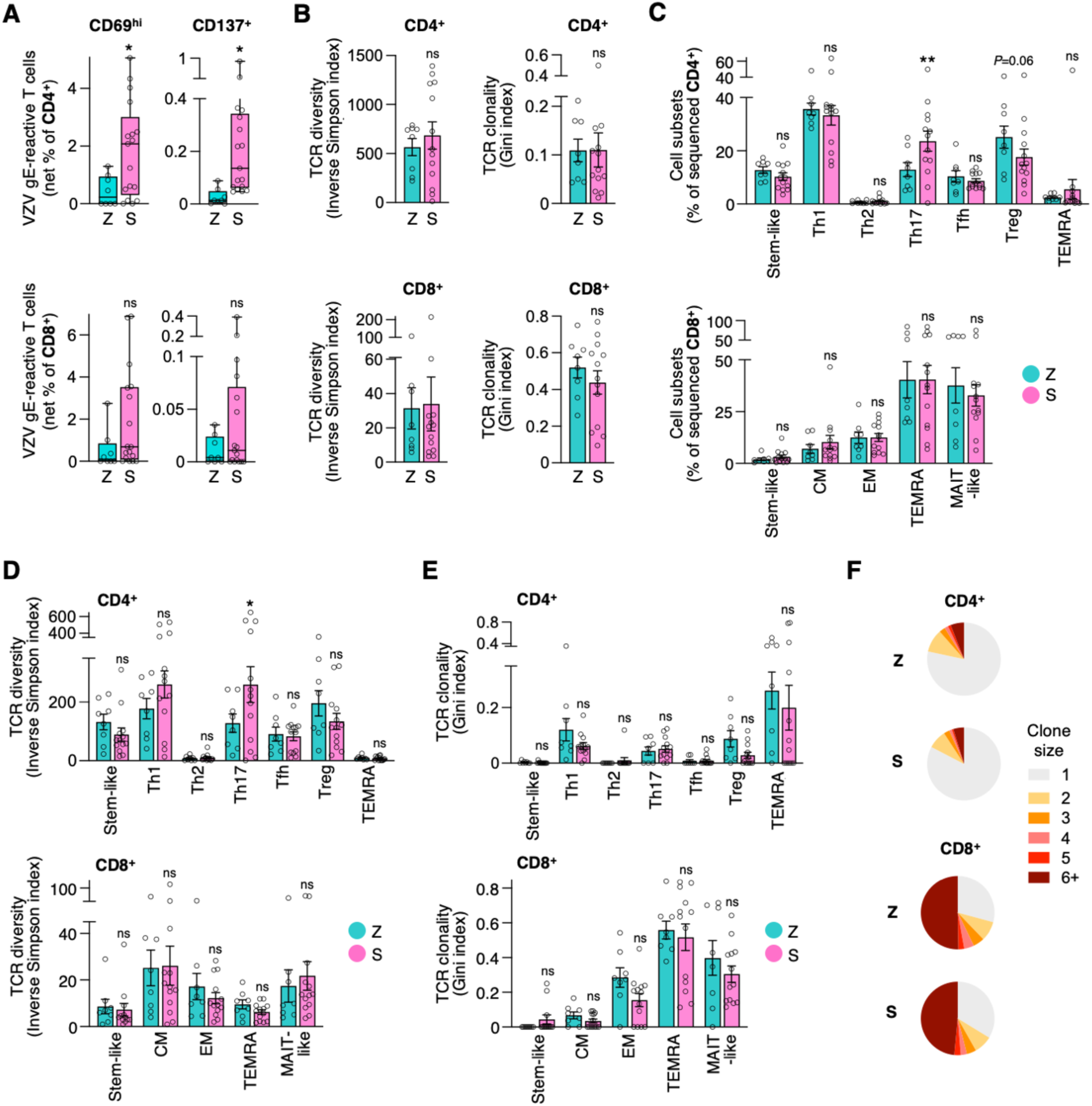
Shingrix vaccination induces an abundant, sustained and Th17-rich T cell response. **A**, Frequencies of VZV gE-reactive CD4^+^ and CD8^+^ T cells from older adults who were vaccinated with the Zostavax (Z) or Shingrix (S) vaccine. VZV gE-reactive T cells were identified as either CD69^hi^ or CD137^+^ by flow cytometry; results are shown as background control-subtracted (net) frequencies. **B**, TCR diversity and clonality indices based on single cell TCRβ CDR3 sequences in VZV gE-reactive CD4^+^ and CD8^+^ T cells from Z and S vaccine recipients. **C**, Distribution of CD4^+^ and CD8^+^ VZV gE-reactive T cells from Z or S vaccine recipients across T cell subsets derived from the single cell RNA-sequencing cluster analysis shown in Supplementary Figure 7. **D-E**, CD4^+^ and CD8^+^ subsets from Z and S recipients are compared for TCR diversity (D) and clonality indices (E). **F**, Pie chart of clonal expansion with size of the pie slices showing the proportion of clones of the indicated size. Data show median (A) or mean ± SEM (B-E). All datapoints represent distinct biological replicates. Data were compared by Mann-Whitney tests (A), two-tailed, unpaired *t*-tests (B-left,C), two-way ANOVA with Šídák’s multiple comparisons test (B-right,D). \**P*<0.05, \*\**P*<0.01. ns, not significant.

When determining whether the longer durability of protective immune memory was associated with T cell subset shifts (Supplementary Figure 7A-G), we did not find any significant differences in probability vectors describing global subset distributions, neither for VZV gE-specific CD4^+^ nor CD8^+^ T cells (Supplementary Figure 7H). However, we noted a significant increase in CD4^+^ Th17 cells as well as a trend towards decreased Tregs in Shingrix recipients (Figure 4C). Increased Th17 cell numbers were associated with higher TCR diversity (Figure 4D) without affecting clonality (Figure 4E,F), possibly indicating recruitment of new T cell specificities upon vaccination into this compartment (*32*). VZV gE-specific CD8^+^ T cells did not differ between Shingrix and Zostavax recipients suggesting that Shingrix vaccination did not simply rejuvenate the VZV gE response to a status seen in young vaccine recipients (Figure 1-2). Rather, Shingrix vaccination appeared to mobilize the CD4^+^ T cell response to confer vaccine protection.

### Plasticity in VZV gE-responsive Th17 T cells in Shingrix and Zostavax recipients

VZV gE-specific CD4^+^ Th17 cells not only differed in frequencies but also showed diverging transcriptomes between Shingrix and Zostavax recipients. PCA of the most variable transcripts showed a trend towards separation of Zostavax and Shingrix recipients for VZV gE-specific CD4^+^ T cells (Supplementary Figure 8A), but not for CD8^+^ T cells (Supplementary Figure 8B). CD4^+^ T cell clusters were unevenly affected by transcript differences with many more DEGs (>1000) for clusters with Th17 and SCM signatures (Figure 5A). Enriched pathways of Shingrix-induced DEGs in SCM cluster 12 included type I and II interferon-related genes (Figure 6B), thereby reversing the age-associated decline in interferon response shown in Figure 3. Notably, this higher expression was predominantly driven by female subjects. For the Th17 clusters C4, C6 and C8, Shingrix recipients showed increased expression of genes indicative of superior functionality (Figure 6C, Supplementary Figure 8C,D). In addition to interferon-related genes, TCR and cytokine signaling, survival and stress-response pathways were enriched. Differential expression of these genes in the Th17 subsets included all recipients irrespective of their gender. GSEA of these Th17 clusters C4, C6, and C8 showed a significant correlation with a Th1 gene signature in Shingrix recipients, suggesting pro-inflammatory potential and possibly polyfunctionality (Figure 5D, Supplementary Figure 8E). In contrast, DEGs with higher expression in VZV gE-responsive Th17 cells of Zostavax recipients were indicative of a Treg signature, with higher expression of classical Treg markers *FOXP3*, *IKZF2*, *CTLA4*, as well as genes promoting Treg stability and function, *RCOR1*(*33*), *KLF2*(*34*), *RBPJ*(*35*), *TOX* and *BATF*(*36, 37*) (Figure 5D,E). These data suggest divergent phenotypic skewing of antigen-specific Th17 cells with Shingrix prioritizing pro-inflammatory, Th1-like features and Zostavax predisposing to immunosuppressive Treg traits.

**Figure 5:**
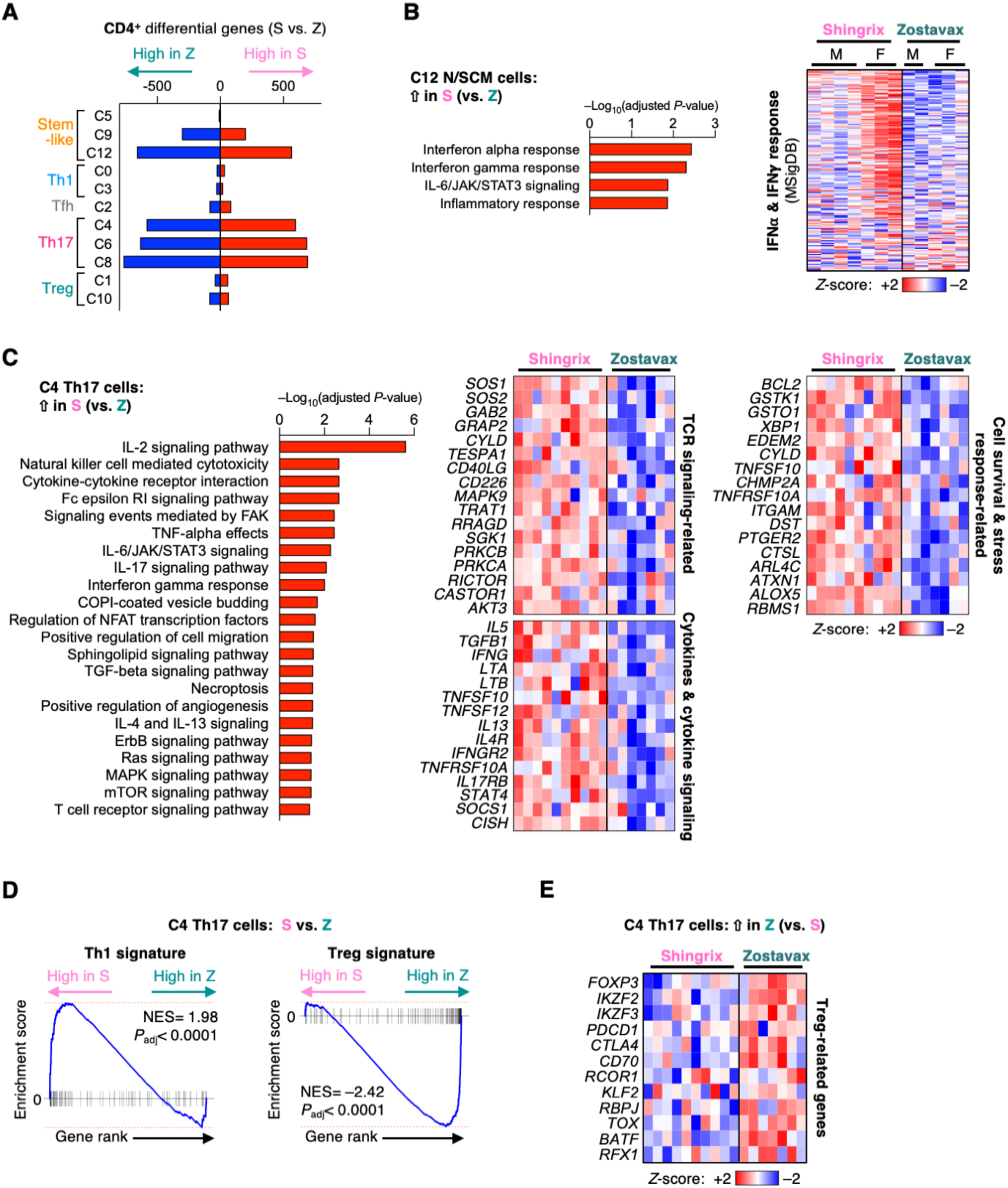
Gene expression in Th17 T cells indicates increased functionality in Shingrix vaccinees. **A**, DEGs in each CD4^+^ scRNA-seq cluster as identified by pseudo-bulk differential gene expression between S and Z vaccine recipients. Only clusters with minimum sample representation of 2 male and 2 female participants per group are included. **B-C**, Pathway enrichment of C12 (B) and C4 (C) DEGs higher expressed in S than in Z (left). Pseudo-bulk gene expression heatmaps of “IFN⍺ response” plus “IFN𝛄 response” gene sets (B, right) and selected DEGs of indicated pathways (C, right). **D**, GSEA of C4 pseudo-bulk gene expression in Z vs S for concordance with Th1 and Treg gene sets. NES, normalized enrichment score. **E**, Pseudo-bulk heatmap of selected DEGs higher expressed in Z as compared to S.

**Figure 6:**
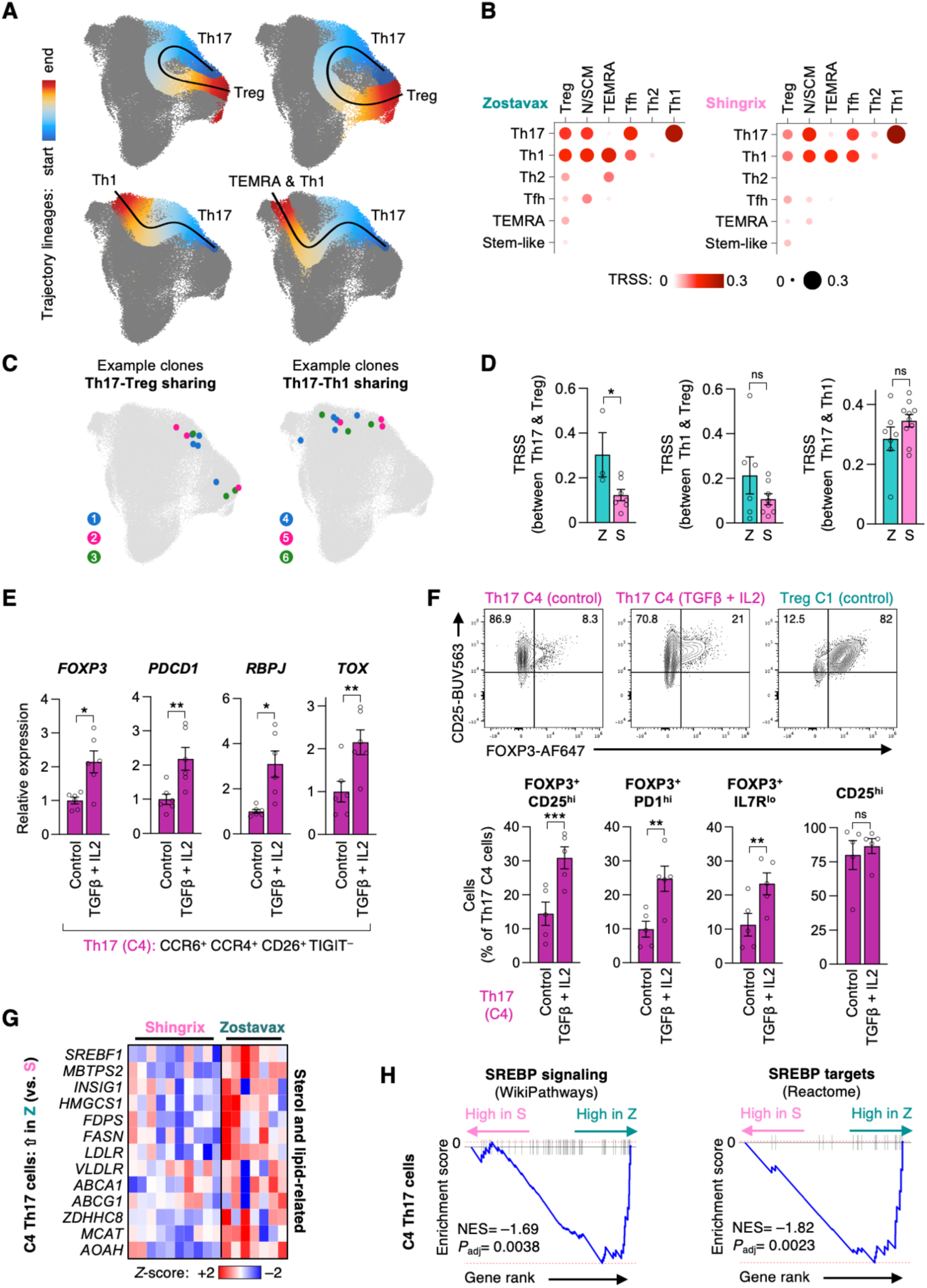
Th17 cells differentiate towards regulatory phenotypes. **A**, Slingshot trajectory analysis projected on the UMAP space. Th17 cluster 4 was selected as starting point to assess putative differentiation potentials. **B**, Bubble plot of clonotype tracking across CD4^+^ VZV gE-reactive T cell subsets. T cell receptor similarity scores (TRSS) are shown as color gradient and bubble size. **C**, Representative TCR-identical clones with divergent transcriptional phenotypes found across different clusters. **D**, TRSS per vaccine recipient showing the extent of TCR sharing between selected clusters. **E-F**, FACS-collected CD4^+^ memory T cells with a cluster 4 Th17 phenotype were stimulated with αCD3/αCD28 in the absence or presence of 25 ng/ml TGFβ and 500 U/ml IL2 for 3 days (E) or 7 days (F). qPCR from 3 experiments (E) and FACS data from 2 experiments (F) are shown. Cluster 1 cells with a Treg phenotype are shown as a comparison. **G**, Heatmap of lipid metabolism-related DEGs in Th17 C4. **H**, GSEA of SREBP (*SREBF1*) signaling and transcriptional target genes in Th17 C4 cells. Data show mean ± SEM (D-F). All datapoints represent distinct biological replicates. Data were compared by two-tailed, unpaired *t*-tests (D) or two-tailed, paired *t*-tests (E,F). \**P*<0.05, \*\**P*<0.01, \*\*\**P*<0.001. ns, not significant.

### Opposing differentiation trajectories of VZV gE-responsive Th17 cells

Th17 cells are capable of phenotypic and functional plasticity along the inflammatory and suppressive T cell spectrum including the differentiation into non-classical Th1* cells(*38*). We therefore investigated the differentiation potential of VZV gE-specific Th17 cells using trajectory analyses on our single cell sequencing data (Figure 6A). Putative trajectories between Th17 C0 cells and either Th1 clusters or regulatory clusters suggest multi-potent differentiation potentials of antigen-specific Th17 cells. We further analyzed TCR clonotype sharing across T cell subsets(*39*) which relies on cells containing the same TCR but divergent gene expression patterns and subset affiliations. TCR sharing was found between the Th17 and Th1 lineages as well as between Th17 and regulatory cells (Figure 6B,C). Interestingly, Zostavax recipients exhibited higher proportions of TCR sharing (∼30%) between Th17 and regulatory cells as compared to ∼10% in Shingrix recipients (Figure 6B,D). In contrast, Th17-Th1 sharing did not differ between vaccine groups. These data suggest that VZV gE-specific Th17 cells have the potential to mis-differentiate into regulatory cells, a process that is more enabled in Zostavax recipient cells.

To experimentally test whether cells with a Th17 (C4) phenotype can acquire features of regulatory cells, we FACS-collected these cells, stimulated them polyclonally in the absence or presence of classical Treg-inducing cytokine combination TGFβ and IL2. After 3 days, qPCR analysis for the Treg marker *FOXP3*, as well as Zostavax Treg-biased Th17 signature genes *PDCD1*, *RBPJ* and *TOX* (Supplementary Figure 9A, see also Figure 5E) confirmed that Th17 cells can acquire Treg traits (Figure 6E). Flow cytometry analysis of these cells after 7 days corroborated the gain of a FOXP3^+^ Treg phenotype in Th17 cells after treatment with TGFβ and IL2 (Figure 6F, Supplementary Figure 9B,C).

We also investigated molecular pathways that may contribute to the differentiation fates of VZV gE-reactive Th17 cells towards regulatory features. Among the top upregulated genes in Zostavax Th17 C4 cells, we noticed *SREBF1*, encoding SREBP1, a master transcription factor of lipogenesis, and many of its associated enzymes and downstream targets including the SREBP- activating enzyme *MBTPS2* encoding S2P, *HMGCS1*, *FPS*, and *FASN* (Figure 7G). GSEA on SREBP signaling and transcriptional target gene sets confirmed higher SREBP activity in Th17 cells of Zostavax individuals (Figure 7H). Lipid metabolic pathways involving fatty acids and sterols, including those controlled by SREBP, have been intricately linked to T cell phenotypes and function(*40*), particularly promoting Treg fitness and suppressiveness(*41, 42*) as well as modulating pro-inflammatory or suppressive Th17 fates(*43, 44*). In summary, we conclude that Th17 cells, particularly those being induced after Zostavax vaccination, can mis-differentiate towards regulatory phenotypes.

## DISCUSSION

Durability of immune memory is of critical importance in designing vaccination strategies and determines the need for booster vaccination. Waning of immune memory is of particular relevance to the older population where it contributes to the increased susceptibility to infections and a poorer vaccine response. In considering mechanisms of durability, it is important to note that memory cells are a dynamic population. Memory is not conferred by the longevity of individual cells but by a population of cells that individually are more short-lived(*45*). Memory T and B cells have limited lifespans, mostly considerably shorter than the duration of immunological memory. The lifespan of a human memory T cell is 30-160 days, in contrast to the typical half-life of human T cell memory of 8-15 years(*46–49*). One exception is a small population of long-lived, stem-like memory cells that have a high degree of self-renewal and a half-life of approximately 9 years(*50*). Memory therefore reflects population averaging of a very diverse set of cells that are subject to selection pressure even in the absence of iterative or chronic antigenic stimulation.

Here, we used VZV vaccination as a model system to uncover determinants of an effective, durable T cell memory by comparing the adaptations of VZV-specific memory T cells many years after vaccination. Specifically, we examined to which extent cell lifespan, contraction in TCR diversity, transition between different functional states and development of cellular senescence or exhaustion contribute to a lack in durable T cell memory. We find that age-associated changes comparing young and older adults vaccinated with a live-attenuated virus vaccine are much more prominent for antigen-specific CD8^+^ than CD4^+^ T cells. The adjuvanted component vaccine Shingrix induces superior durability without restoring these defects in CD8^+^ immunity but by inducing compensatory properties in CD4^+^ T cells. Moreover, neither VZV gE- responsive CD4^+^ nor CD8^+^ T cells acquire a transcriptional signature of cellular senescence or exhaustion. This is consistent with a recent study in which antigen-specific CD8^+^ T cells in mice undergoing iterative viral infections have an infinite lifespan as long as there is a minimal interval of several weeks between stimulations(*51*).

An essential role in maintaining long-term memory, in particular for CD8^+^ T cells, has been attributed to a small population of stem-like memory T cells that shares features of naïve and memory cells and that is able to reconstitute the full spectrum of memory and effector T subpopulations(*50*). Transcription factors that are pivotal for preserving stemness include TCF1 and LEF1. A decline in TCF1 levels is a hallmark of the aging process in global naïve CD4^+^ T cells(*52*), and lower expression of TCF1 in CD8^+^ than CD4^+^ naïve T cells may contribute to their higher loss with aging. Frequencies of VZV gE-reactive SCM trended to be reduced with age for both CD4^+^ and CD8^+^ T cells and DEGs in the SCM subsets included several genes of functional importance to stemness, suggesting that this mechanism contributed to the loss in vaccine protection. However, vaccination with Shingrix did not reverse these defects demonstrating that the superior memory durability was not directly related to improved SCM.

Immune protection depends on a diverse TCR repertoire. Both Zostavax and Shingrix vaccination can recruit new CD4^+^ T cell specificities into the memory compartment, thereby diversifying the repertoire(*32, 53*). Obviously, this depends on a sizeable naïve compartment, which is largely preserved with aging in CD4^+^ but much less so for CD8^+^ T cells. In addition, the repertoire of memory T cells is under selection pressures, in particular in a setting of latent infection such as VZV. We found VZV gE-specific CD4^+^ T cells to be highly diverse across different subsets except TEMRA cells, with only marginal clonality increase in older vaccine recipients. In contrast, VZV gE-specific CD8^+^ T cells were less diverse compared to CD4^+^ T cells already in young adults, consisting with previous studies on the global memory repertoire(*54*). Moreover, they lose diversity with age. Although Shingrix’ adjuvants monophosphoryl lipid A (MPL) and QS-21 can also stimulate CD8^+^ T cell responses(*31, 55*), Shingrix vaccination did not restore the age-dependent decline in CD8^+^ T diversity, possibly due to older adults lacking the source of naïve CD8^+^ T cells. Contraction in CD8^+^ T cell diversity may therefore explain the decline in VZV vaccine protection with age, while it does not explain the superior protection by Shingrix.

Instead, Shingrix vaccination prioritizes the generation of Th17 CD4^+^ T cells while tending to evade Treg phenotypes. Shingrix likely induces Th17 generation through the AS01_B_ adjuvant system, which is also largely responsible for the superior vaccine protection(*56*). AS01_B_ includes the saponin QS-21 and the TLR4 agonist MPL. TLR4 activation in CD4^+^ T cells indeed promotes Th17 responses in the experimental autoimmune encephalomyelitis (EAE) mouse model(*57*). Additionally, TLR4 activation in antigen-presenting cells leads to the production of inflammatory cytokines including IL6, IL1β, IL12, and TNFα, with the potential to skew the milieu for CD4^+^ T helper differentiation from Treg- to Th17-promoting conditions(*58, 59*). Th17 responses have been associated with effective vaccine responses in other contexts(*60*), such as vaccination with the acellular pertussis vaccine(*16, 17, 61*), the adjuvanted VP6 component rotavirus vaccine in mice(*60*), or the Th17-skewing, autologous dendritic cell vaccines for ovarian cancer patients(*62*).

Our data also indicate that Th17 heterogeneity and plasticity distinguishes durable from short-lived VZV immune memory in older adults. Th17 cells that are prone to acquire Treg features appear less protective than those that are poised towards pro-inflammatory Th1 phenotypes. This Th17 phenotypic spectrum resembles the one previously described for EAE with Treg-like phenotypes being consistent with homeostatic Th17 cells, while Th1-like phenotypes matching those of EAE-pathogenic Th17 cells(*38, 44*). In line with previous reports, we find that lipid metabolic signaling associated with adapting these opposing Th17 fates(*40*). More broadly, differentiation of Th17 into classical Treg or Tr1 cells has been described by fate mapping in mice(*39, 63, 64*). Unwarranted mis-differentiation of VZV gE-reactive Th17 cells into Tregs may therefore contribute to the higher accumulation of antigen-reactive Tregs in Zostavax recipients and, thus, to the lower vaccine efficacy and durability of Zostavax. Our observations are supported by our TCR clonotype sharing data, and by the fact that virus antigen-specific Treg cells have been described in relatively high proportions in human blood and in multiple contexts such as HIV, HSV, or CMV(*65*) as well as in the skin after VZV challenge(*66*). Thus, the higher Th17 numbers combined with their pro-inflammatory, Th1-like character in the Shingrix response are associated with superior and durable vaccine protection in older adults.

Our studies have limitations inherent to studying human immunology in a diverse population. The VZV vaccination system is well suited to study maintenance of immune memory with antigen-specific T cells being protective rather than humoral immunity. We focused on the gE antigen that is shared between the three vaccines, but the immune response is broader for the live-attenuated VZV vaccine (*67*). However, the interpretation of read-out systems used here such as repertoire diversity or differential gene expression should not be affected by this difference. Our post-vaccination time intervals differed for Zostavax and Shingrix, given that Zostavax was discontinued shortly after Shingrix was introduced. It can therefore not completely be excluded that Shingrix recipients eventually develop a similar misdifferentiation phenotype in antigen-specific Th17 cells at later timepoints. Our study populations were small, in part due to the declining number of individuals that had a Zostavax vaccination in recent years. We included male and female participants but populations were too small to follow up on observations of a gender-specific effect. Finally, given the substantial heterogeneity of MHC polymorphism in human populations, we had to rely on a functional definition of antigen-specificity based on the expression of activation markers. Nonetheless, our study indicates the potential of leveraging Th17 CD4^+^ memory T cell responses in vaccination strategies for older adults. Inclusion of Th17- inducing adjuvants, such as AS01_B_, in vaccines would be a relevant area for future research and development.

## MATERIALS AND METHODS

### Study design and study populations

In this study, we sought to determine the molecular determinants of T cell memory durability in older adults and employed VZV vaccination as a model system. We contrasted VZV gE-specific T cells derived from peripheral blood of (i) young adults (<30 years) who received a childhood live-attenuated VZV vaccine, Varivax, which elicits long lasting immune memory and protection against VZV; (ii) older adults (>55 years) who received the same live-attenuated vaccine strain, marketed as Zostavax, which induces a short-term, partial protection against VZV reactivation; and (iii) older adults (>55 years) who received an effective, durable adjuvanted vaccine, Shingrix. We recruited a total of 53 volunteers who did not have an acute or active chronic disease, cancer or autoimmune disease, and were not on a chronic anti-inflammatory medication and for whom the vaccination history was documented. Recruitment was done by the Mayo Clinic Biobank registry. The sample size of Zostavax recipients was restricted by the number of individuals in the registry who fulfilled the inclusion criteria and agreed to participate. A similar number of Shingrix recipients were chosen who had received the vaccine early after it became available. We received peripheral blood of these volunteers and isolated peripheral blood mononuclear cells (PBMC). Participants gave informed written consent. Chronic diseases were permitted if controlled by medication. Phlebotomy was scheduled in the morning, and volunteers were asked to fast before the blood collection. Volunteers for single cell sequencing were recruited in 10 batches between December 2021 and July 2023. Subject information on samples used in single cell sequencing experiments is summarized in Supplementary Table 1 and 2. In addition, PBMCs from leukoreduction system chambers of 37 blood donors were purchased from the Mayo Clinic Blood Donor Center, Department of Laboratory Medicine and Pathology - Component Laboratory. Samples were de-identified except for age and self-assigned sex information. These samples were only used for mechanistic studies. Samples from male and female individuals were used. Young adults were 20 to 35 years old, older adults were 60 years or older. Studies involving human subjects were approved by the Mayo Clinic Institutional Review Board.

### Cell isolation, culture, and antigen stimulation

Lymphoprep (StemCell Technologies, #07861) was used to purify PBMCs by density centrifugation according to the manufacturer’s instructions. Cells were used fresh in all experiments and cultured in RPMI 1640 medium supplemented with 5% filtered human AB serum (Sigma Aldrich, #H4522), 100 U/mL penicillin, and 100 U/mL streptomycin (Sigma Aldrich, #P0781) in an incubator with 5% CO_2_ and atmospheric O_2_. For peptide stimulation experiments, we employed activation-induced marker (AIM) assays(*29*). Briefly, PBMCs were seeded at 1x10^6^ cells per U-bottom 96-well (for flow cytometry analyses) or 20x10^6^ per 12-well (for single cell sequencing) and stimulated with 1 μg/mL of a VZV glycoprotein E (gE) peptide pool diluted in 0.2% DMSO (15-mer peptides with 11 amino acids overlap, total of 153 peptides; # PM-VZV-gE, JPT Peptide Technologies). Ultra-LEAF anti-CD28 antibody was added at 1 μg/mL (clone 28.2, BioLegend, #302943). For samples that were subjected to single cell sequencing or flow cytometry to measure CD40L levels, anti-CD40 blocking antibody was added at 1 μg/mL (clone HB14, Miltenyi, #130-094-133). Parallel control PBMC cultures of the same individuals received 0.2% DMSO instead of the VZV gE peptide mix, or 1 μg/mL SEB (Staphylococcal enterotoxin B, Toxin Technologies, #BT202red). PBMCs were spun in plates at 300×g for 3 min before culturing for 42. To calculate net frequences, the proportion of activated T cells in the solvent control cultures was subtracted from the proportion of antigen-responsive T cells. Quantitation and phenotyping of VZV gE-reactive T cells after 42 hours of VZV gE stimulation was performed via cell surface antibody staining or intracellular staining. Utilized antibodies are listed in Supplementary Table 3. Viability dye Live/Dead Fixable Blue (Invitrogen, #L23105) was included. For intracellular staining, cells were fixed with the eBioscience Foxp3/Transcription Factor Staining Buffer Set (Thermo Fisher Scientific, #00-5523-00) according to the manufacturer’s instructions. Most experiments were analyzed on a 5-laser Cytek Aurora (Cytek Biosciences) running SpectroFlo v3 with UltraComp eBeads compensation beads (Thermo Fisher Scientific, #01-2222-42). Only AIM assay optimization experiments were analyzed on a BD LSRFortessa X- 20.

### Statistical analyses

For statistical association of cluster proportions with vaccine groups, we performed probability vector analyses and permutation tests as described previously(*68*). Briefly, each sample is characterized as a probability vector whose components are the relative cell frequency in each of the clusters and the cardinality represents the number of distinct clusters. We performed linear regression analyses with sex adjustment between the individual components of the probability vector to examine the association between the probability vector and vaccine groups. The association with the entire probability vector is summarized by the sum of squares of all *Z*-scores corresponding to the regression coefficient of the vaccine groups from the linear regression analyses and tested via permutation test permuting the vaccine groups.

All other statistical analyses were performed via Prism software 10 (GraphPad). Data are presented as mean with error bars indicating standard error of the mean (SEM), unless otherwise indicated. Unpaired or paired, two-tailed Student’s t-tests were used when comparing two groups. One-way or two-way ANOVA with multiple comparisons test were used for multigroup comparisons. A *P*-value less than 0.05 was considered statistically significant. Significance levels (\**P*< 0.05, \*\**P*<0.01, \*\*\**P*<0.001, \*\*\*\**P*<0.0001) are indicated in figures.

## Data availability

Single cell sequencing data have been deposited at the Gene Expression Omnibus (GSE286045) and are publicly available as of the date of publication.

## ACKNOWLEDGEMENTS

We thank the Mayo Clinic Center for Individualized Medicine, Janet Olson, Nicole Larson, and Bridget Rathbun for their help in participant recruitment, William Greenleaf and Rohit Jadhav for discussions on bioinformatic analyses, Lu Tian for statistical analysis consultation, Yong Han, Drew Kluge, Colleen Moe and Ryan Fallon for cell sorting, and Vernadette Simon and Fred Rakhshan for single cell capture and library preparation. This study was supported with resources and the use of facilities at the Mayo Clinic Microscopy and Cell Analysis Core and Mayo Clinic Medical Genome Facility.

## Autor contributions

I.S., A.J., C.M.W., and J.J.G. conceptualized the study. I.S. performed experiments and analyzed data with help of J.J., H.O., Y.M., and M.O.. A.J. performed bioinformatic analyses. I.S., A.J., and J.J.G. interpreted data and wrote the manuscript. All authors edited the manuscript.

**Please see the Supplementary Materials and Methods for additional method descriptions.**

## LIST OF SUPPLEMENTARY MATERIALS

The Supplementary Materials contains Supplementary Materials and Methods, 9 Supplementary Figures and 5 Supplementary Tables. It also contains 2 figures on flow cytometry gating strategies. Correspondence and requests for materials should be addressed to Jörg. J. Goronzy.

**Supplementary Figure 1:** Young and older vaccine recipients have similar frequencies of VZV gE-reactive T cells.

**Supplementary Figure 2:** CD4^+^ memory T cells specific for VZV gE are phenotypically highly diverse.

**Supplementary Figure 3:** The VZV gE-reactive CD8^+^ memory T cell response includes diverse subsets.

**Supplementary Figure 4:** Stem-like features in the CD4^+^ T cell memory response from older adults against VZV gE are diminished.

**Supplementary Figure 5:** CD8^+^ memory T cells specific to VZV gE lack signatures of age-related exhaustion or cellular senescence.

**Supplementary Figure 6:** VZV gE responses are increased in Shingrix recipients.

**Supplementary Figure 7:** VZV gE-specific T cells in Shingrix and Zostavax vaccine recipients include diverse subsets.

**Supplementary Figure 8:** VZV gE-reactive CD4^+^ T cells in all Th17-related clusters share gene expression profiles indicating increased functionality in Shingrix recipients.

**Supplementary Figure 9:** Signatures of Th17 cells overlap with cluster 1 Treg phenotypes under selective conditions.

**Supplementary Table 1:** Patient demographics for single cell sequencing samples.

**Supplementary Table 2:** Patient comorbidities and key medication for single cell sequencing samples.

**Supplementary Table 3 (Excel):** Sample information for single cell sequencing.

**Supplementary Table 4 (Excel):** Antibodies for Flow Cytometry.

**Supplementary Table 5 (Excel):** TotalSeq-B antibodies for CITE-seq.

**Flow Cytometry Gating Strategy 1:** FACS gating strategy to collect VZV gE-reactive T cells for single cell sequencing.

**Flow Cytometry Gating Strategy 2:** FACS gating strategy to collect phenotypic subsets of VZV gE-reactive T cells for functional studies.

