## Supplementary Information for "Antigen-specific Th17 T cells offset the age-related decline in durable T cell immunity"

**This PFD includes:**

Supplementary Materials and Methods

Supplementary Figures 1-9

Supplementary Tables 1-2

Flow Cytometry Gating Strategy 1-2

**Other Supplementary Information for this manuscript include:**

Supplementary Tables 3-5 (Excel format)

### SUPPLEMENTARY MATERIALS AND METHODS

#### Cell preparation for single cell sequencing

For single cell sequencing experiments, 42 hours after peptide stimulation, cultures were pre-purified with EasySep Human T Cell Isolation Kit (using 20% of the recommended reagents, Stemcell Technologies, # 17951). T cells were counted and resuspended at 1-2 x$10^6$ cells / 50 μL and blocked with TruStain FcX Fc Blocking reagent (BioLegend, #422301) for 10 min at 4ºC before incubation with hashtag TotalSeq-B antibodies (0.5 μL per 100 μL cell suspension, Supplementary Table 4) and flow cytometry antibodies (Supplementary Table 3, and Live/Dead Fixable Aqua, Invitrogen, #L34957) for 30 minutes at 4ºC. Cells were washed once before FACS on BD FACSAria 4-laser digital flow cytometer with FACSDiva v8 software. Cell sorting was performed by the Mayo Clinic Microscopy and Cell Analysis Core Flow Cytometry Lab. Live, single $CD3^+$ $CD8^+$ or $CD3^+$ $CD4^+$ cells that were $CD69^{hi}$ and/or $CD137^+$ were collected. A representative gating strategy is shown in the Supplementary Materials. After sorting, cells were counted and pooled into one sample across T cell subsets and donors at similar cell numbers: VZV gE-responsive $CD4^+$ T cells, DMSO background-activated $CD4^+$ T cells, VZV gE-responsive $CD8^+$ T cells (at about half the proportion of VZV gE-responsive $CD4^+$ T cells), DMSO background-activated $CD8^+$ T cells. The pooled sample was then stained with diluted TotalSeq-B cell surface marker antibodies (BioLegend, Supplementary Table 4) in PBS containing 2% FBS for 30 minutes at 4°C. Cells were washed twice with PBS containing 2% FBS, and cell viability was determined with Trypan Blue which was consistently >90%. Cells were subjected to single cell capture via the Chromium controller (10x genomics), Chromium Next GEM Single Cell 3' Kit v3.1 (Single Index, 10x genomics, #PN-1000121) and gel beads (10x genomics, #1000376). About 20,000 cells per pooled samples were targeted for cell capture. Three libraries were generated from each captured sample: 3' gene expression library, library of antibody-derived tags (ADT) for TotalSeq-B antibodies including hashtag antibodies and cell surface marker (3' Feature Barcode Library Kit,

10x genomics, #PN-1000079) collectively known as CITE-seq(*1*). The third library was constructed to capture TCR CDR3 sequences via a custom protocol (see section below).

**Functional testing of VZV gE-reactive T cell subsets**

PBMCs were stimulated with VZV gE peptides for 42 hours as above. $CD4^+$ T cells were purified from these cultures via the EasySep Human $CD4^+$ T Cell Isolation Kit (Stemcell Technologies, #17952 using 20% of the recommended reagents). Cells were stained with flow cytometry antibodies and VZV gE-reactive T cell subsets were FACS-collected via a BD FACSAria. Cells were pre-gated to be live, single $CD3^+$ $CD4^+$ cells that were $CD69^{hi}$ and/or $CD137^+$ and further subdivided via single cell cluster-defining markers: $CCR6^+$ $CCR4^-$ $IL7R^+$ (cluster 0/Th1), $CCR6^+$ $CCR4^+$ $TIGIT^+$ (cluster 1/Treg), $CCR6^+$ $CCR4^+$ $CD26^+$ $TIGIT^-$ (cluster 4/Th17). A representative gating strategy is shown in the Supplementary Materials. Sorted cells were stained with 2 μM CellTrace Violet (Invitrogen, #C34557) according to the manufacturer's instructions. Cells were counted and seeded into 96-well plates (100,000 cells/well, 100 μL volume) that had been coated with αCD3/αCD28 antibodies (both 1 μg/mL; clones UCHT1 and CD28.8, respectively; BioLegend, #300465 and #302943). After 5 days of culture, the culture supernatant was collected and frozen at –80°C. Cytokine levels in these culture supernatants were measured with LEGENDPlex 12-plex Human Th Cytokine Panel (BioLegend, # 741028) according to the manufacturer's instructions. Cells were also collected, stained with flow cytometry antibodies (Supplementary Table 3), viability dye Live/Dead Fixable Blue (Invitrogen, #L23105) and ApoTracker Green (BioLegend, #4274020). Samples were analyzed on a 5-laser Cytek Aurora (Cytek Biosciences). UltraComp eBeads compensation beads (Thermo Fisher Scientific, #01-2222-42) were used for single staining controls.

**Single cell sequencing and data processing**

Single cell gene expression and ADT libraries were sequenced on a NovaSeq 6000 S4 (Illumina) and 150 cycle, paired-end kit to a target depth of 50,000 read pairs/cell for gene expression, 5,000 read pairs/cell for feature barcodes of the ADT/CITE libraries. Cell capture and library construction was performed by the Mayo Clinic Medical Genome Facility Genome Analysis Core and sequencing was performed by Novogene Corporation or Azenta Life Sciences. Data were similarly processed as described previously. Fastq reads were assessed using FastQC v0.11.9. Following quality assessment, raw reads were aligned to the GRCh38 human genome using Cell Ranger multi v7.1.0(*2*). The Cell Ranger multi pipeline was employed to quantify libraries of antibody-derived tags (ADT) and demultiplex hash-tagged samples of individual donors in pooled libraries. Seurat v4.2.1 was used for downstream analysis following data preprocessing(*3, 4*). Quality control was conducted to exclude low-quality cells based on the following filters: cells with <200 or >5000 expressed genes, >10% UMIs mapping to mitochondria-encoded transcripts, or >20,000 ADT UMIs. DoubletFinder was applied to identify and remove cell doublets(*5*). We identified $CD4^+$ versus $CD8^+$ T cells based on their CD4 and CD8$\alpha$ ADT protein as well as *CD4* and *CD8A* gene expression. $CD4^+$ T cells were CD4 ADT-positive but CD8$\alpha$ ADT-negative, or *CD4* RNA-expressing but *CD8A* RNA-negative. $CD8^+$ T cells were CD8$\alpha$ ADT-positive but CD4 ADT-negative, or *CD8A* RNA-expressing but *CD4* RNA-negative. Across all experiments and groups, we obtained 137,461 cells of which 97,663 were $CD4^+$ T cells and 39,798 were $CD8^+$ T cells. $CD4^+$ and $CD8^+$ T cells were further processed separately. For gene expression normalization, individual samples were processed using SCTransform(*6*) while regressing out mitochondrial gene expression. ADT counts were normalized using centralized log-ratio (CLR) transformation. Batches across experiments were integrated using diagonalized Canonical Correlation Analysis (CCA) to identify mutual nearest neighbors that act as anchors employing the FindIntegrationAnchors and IntegrateData functions(*4*). The integrated gene expression and

ADT modalities, respective matrices were scaled, and principal components were calculated. A weighted nearest-neighbor (WNN) graph was constructed using the FindMultiModalNeighbors function(*3*) and used to generate the UMAP embeddings, facilitating the visualization for both the modalities. Clustering was performed based on WNN using smart local moving algorithm (SLM)(*3*). We performed imputation of gene expression using Markov Affinity-based Graph Imputation of Cells (MAGIC) solely for data visualization(*7*). To assist in annotations of clusters to classical T cell subsets, we performed UCell gene module scores using the AddModuleScore_UCell function with published T cell gene signatures(*8, 9*). Pseudotime trajectory analysis was performed with SlingShot v.2.14.0(*10*) using $CD4^+$ T cell cluster C4 as starting point.

**Pseudobulk differential expression analysis**

Raw counts of samples were aggregated by summing the UMIs from their respective single cells utilizing Seurat's built-in function AggregateExpression(*3*). Samples containing fewer than 20 cells were excluded from this analysis. Low-expressing genes with fewer than 10 counts in the selected comparisons were removed using the filterByExpr function(*11*). The remaining raw counts were transformed using regularized log (rlog) transformation to compute principal components, followed by batch effect correction with plotPCA and removeBatchEffect functions, respectively(*11, 12*). Differentially expressed genes (DEGs) were identified by comparing age and vaccine groups across all cells, within a given cluster as well as between the clusters. Only clusters with minimum sample representation of 2 male and 2 female participants per group were included. DEGs were identified by modeling the mean-variance trend on log-transformed counts per million (logCPM) of each gene using voomWithQualityWeights from the limma package(*12, 13*). For Shingrix and Zostavax comparison the experimental batch was incorporated as a covariate. Contrasts were fitted to the model, and empirical Bayes moderation was applied to identify DEGs with an adjusted *P*-value < 0.05. Pathway enrichment analyses on DEGs were conducted with EnrichR(*14*) focusing on Gene Ontologies, Reactome, BioPlanet, KEGG, MSigDB, and WikiPathways gene sets. Gene set enrichment analysis (GSEA) was performed by FGSEA v1.20.0[(15)] for adaptiveness and lymphocyte innateness genes(*16*), exhaustion-associated genes (GSE9650, GSE41867), cellular senescence-associated genes (Reactome R-HSA-2559583, SenMayo[(17)]), T helper signatures(*8, 9*), SREBP-related genes (Reactome R-HSA-2426168, WikiPathways WP1982).

**Single cell T cell receptor sequencing and data analysis**

Single cell libraries for TCR sequences were created from the same 3' single cell, full-length cDNA material used for gene expression and ADT libraries. *TRB* libraries were generated as described previously(*18-20*). For one experiment, we also generated *TRA* libraries. Libraries were sequenced on MiSeq or NextSeq 2000 sequencers (Illumina) with a 100-cycle single-end kit with TruSeq Read 1 primer and custom index 1 primer (cycle 28-100-0-0). We targeted at least 2,500 read pairs/cell. Fastq files were processed with WAT3R(*21*) to identify the TCR chains and CDR3 sequences. We obtained a *TRB* CDR3 sequence from 38% of $CD4^+$ or $CD8^+$ T cells that were included in our transcriptomic analyses. We calculated the Inverse Simpson index to estimate TCRβ diversity and the Gini index to calculate TCRβ clonality as described previously(*22*). For clonotype tracking across T cell subsets we calculated the TCR repertoire similarity score (TRSS)(*23*).

**RNA isolation and qRT-PCR**

T cells were washed with PBS, the cell pellet was lysed in RLT buffer and RNA was isolated with the RNeasy Micro kit (Qiagen, #74004) according to the manufacturer's instructions but avoiding the DNase digestion step. About 200 ng RNA was used for reverse transcription via the SuperScript VILO cDNA Synthesis Kit (Invitrogen, #11754050). The PowerUp SYBR Green Master Mix (Applied Biosystems, #A25742) was used for qPCR analyses on a QuantStudio 6 (Applied Biosystems). Primers were: *FOXP3* forward 5'-GAAACAGCACATTCCCAGAGTTC-3', *FOXP3* reverse 5'-ATGGCCCAGCGGATGAG -3', *PDCD1* forward 5'-AAGGCGCAGATCAAAGAGAGCC-3', *PDCD1* reverse 5'-CAACCACCAGGGTTTGGAACTG-3', *RBPJ* forward 5'-TCATGCCAGTTCACAGCAGTGG-3', *RBPJ* reverse 5'-TGGATGTAGCCATCTCGGACTG-3', *TOX* forward 5'-CGCTACCTTTGGCGAAGTCTCT-3', *TOX* reverse 5'-CTGGCTCTGTATGCTGCGAGTT-3'. *ACTB* was used as reference gene, forward 5'-CACCATTGGCAATGAGCGGTTC-3' , reverse 5'-AGGTCTTTGCGGATGTCCACGT-3'.

### REFERENCES AND NOTES (for the Supplementary Materials and Methods)


1. M. Stoeckius, C. Hafemeister, W. Stephenson, B. Houck-Loomis, P. K. Chattopadhyay, H. Swerdlow, R. Satija, P. Smibert, Simultaneous epitope and transcriptome measurement in single cells. *Nat Methods* **14**, 865-868 (2017).
2. G. X. Zheng, J. M. Terry, P. Belgrader, P. Ryvkin, Z. W. Bent, R. Wilson, S. B. Ziraldo, T. D. Wheeler, G. P. McDermott, J. Zhu, M. T. Gregory, J. Shuga, L. Montesclaros, J. G. Underwood, D. A. Masquelier, S. Y. Nishimura, M. Schnall-Levin, P. W. Wyatt, C. M. Hindson, R. Bharadwaj, A. Wong, K. D. Ness, L. W. Beppu, H. J. Deeg, C. McFarland, K. R. Loeb, W. J. Valente, N. G. Ericson, E. A. Stevens, J. P. Radich, T. S. Mikkelsen, B. J. Hindson, J. H. Bielas, Massively parallel digital transcriptional profiling of single cells. *Nat Commun* **8**, 14049 (2017).
3. Y. Hao, S. Hao, E. Andersen-Nissen, W. M. Mauck, 3rd, S. Zheng, A. Butler, M. J. Lee, A. J. Wilk, C. Darby, M. Zager, P. Hoffman, M. Stoeckius, E. Papalexi, E. P. Mimitou, J. Jain, A. Srivastava, T. Stuart, L. M. Fleming, B. Yeung, A. J. Rogers, J. M. McElrath, C. A. Blish, R. Gottardo, P. Smibert, R. Satija, Integrated analysis of multimodal single-cell data. *Cell* **184**, 3573-3587 e3529 (2021).
4. T. Stuart, A. Butler, P. Hoffman, C. Hafemeister, E. Papalexi, W. M. Mauck, 3rd, Y. Hao, M. Stoeckius, P. Smibert, R. Satija, Comprehensive Integration of Single-Cell Data. *Cell* **177**, 1888-1902 e1821 (2019).
5. C. S. McGinnis, L. M. Murrow, Z. J. Gartner, DoubletFinder: Doublet Detection in Single-Cell RNA Sequencing Data Using Artificial Nearest Neighbors. *Cell Syst* **8**, 329-337 e324 (2019).
6. C. Hafemeister, R. Satija, Normalization and variance stabilization of single-cell RNA-seq data using regularized negative binomial regression. *Genome Biol* **20**, 296 (2019).
7. D. van Dijk, R. Sharma, J. Nainys, K. Yim, P. Kathail, A. J. Carr, C. Burdziak, K. R. Moon, C. L. Chaffer, D. Pattabiraman, B. Bierie, L. Mazutis, G. Wolf, S. Krishnaswamy, D. Pe'er,

Recovering Gene Interactions from Single-Cell Data Using Data Diffusion. *Cell* **174**, 716-729 e727 (2018).
8. B. J. Meckiff, C. Ramirez-Suastegui, V. Fajardo, S. J. Chee, A. Kusnadi, H. Simon, S. Eschweiler, A. Grifoni, E. Pelosi, D. Weiskopf, A. Sette, F. Ay, G. Seumois, C. H. Ottensmeier, P. Vijayanand, Imbalance of Regulatory and Cytotoxic SARS-CoV-2-Reactive CD4(+) T Cells in COVID-19. *Cell* **183**, 1340-1353 e1316 (2020).
9. G. Seumois, C. Ramirez-Suastegui, B. J. Schmiedel, S. Liang, B. Peters, A. Sette, P. Vijayanand, Single-cell transcriptomic analysis of allergen-specific T cells in allergy and asthma. *Sci Immunol* **5**, (2020).
10. K. Street, D. Risso, R. B. Fletcher, D. Das, J. Ngai, N. Yosef, E. Purdom, S. Dudoit, Slingshot: cell lineage and pseudotime inference for single-cell transcriptomics. *BMC Genomics* **19**, 477 (2018).
11. M. D. Robinson, D. J. McCarthy, G. K. Smyth, edgeR: a Bioconductor package for differential expression analysis of digital gene expression data. *Bioinformatics* **26**, 139-140 (2010).
12. M. E. Ritchie, B. Phipson, D. Wu, Y. Hu, C. W. Law, W. Shi, G. K. Smyth, limma powers differential expression analyses for RNA-sequencing and microarray studies. *Nucleic Acids Res* **43**, e47 (2015).
13. R. Liu, A. Z. Holik, S. Su, N. Jansz, K. Chen, H. S. Leong, M. E. Blewitt, M. L. Asselin-Labat, G. K. Smyth, M. E. Ritchie, Why weight? Modelling sample and observational level variability improves power in RNA-seq analyses. *Nucleic Acids Res* **43**, e97 (2015).
14. M. V. Kuleshov, M. R. Jones, A. D. Rouillard, N. F. Fernandez, Q. Duan, Z. Wang, S. Koplev, S. L. Jenkins, K. M. Jagodnik, A. Lachmann, M. G. McDermott, C. D. Monteiro, G. W. Gundersen, A. Ma'ayan, Enrichr: a comprehensive gene set enrichment analysis web server 2016 update. *Nucleic Acids Res* **44**, W90-97 (2016).
15. G. Korotkevich, V. Sukhov, N. Budin, B. Shpak, M. N. Artyomov, A. Sergushichev, Fast gene set enrichment analysis. *BioRxiv*, (2021).
16. M. Gutierrez-Arcelus, N. Teslovich, A. R. Mola, R. B. Polidoro, A. Nathan, H. Kim, S. Hannes, K. Slowikowski, G. F. M. Watts, I. Korsunsky, M. B. Brenner, S. Raychaudhuri, P. J. Brennan, Lymphocyte innateness defined by transcriptional states reflects a balance between proliferation and effector functions. *Nat Commun* **10**, 687 (2019).
17. D. Saul, R. L. Kosinsky, E. J. Atkinson, M. L. Doolittle, X. Zhang, N. K. LeBrasseur, R. J. Pignolo, P. D. Robbins, L. J. Niedernhofer, Y. Ikeno, D. Jurk, J. F. Passos, L. J. Hickson, A. Xue, D. G. Monroe, T. Tchkonia, J. L. Kirkland, J. N. Farr, S. Khosla, A new gene set identifies senescent cells and predicts senescence-associated pathways across tissues. *Nat Commun* **13**, 4827 (2022).
18. T. E. Miller, C. A. Lareau, J. A. Verga, E. A. K. DePasquale, V. Liu, D. Ssozi, K. Sandor, Y. Yin, L. S. Ludwig, C. A. El Farran, D. M. Morgan, A. T. Satpathy, G. K. Griffin, A. A. Lane, J. C. Love, B. E. Bernstein, V. G. Sankaran, P. van Galen, Mitochondrial variant enrichment from high-throughput single-cell RNA sequencing resolves clonal populations. *Nat Biotechnol* **40**, 1030-1034 (2022).
19. A. A. Tu, T. M. Gierahn, B. Monian, D. M. Morgan, N. K. Mehta, B. Ruiter, W. G. Shreffler, A. K. Shalek, J. C. Love, TCR sequencing paired with massively parallel 3' RNA-seq reveals clonotypic T cell signatures. *Nat Immunol* **20**, 1692-1699 (2019).
20. Y. Sato, A. Jain, S. Ohtsuki, H. Okuyama, I. Sturmlechner, Y. Takashima, K. C. Le, M. C. Bois, G. J. Berry, K. J. Warrington, J. J. Goronzy, C. M. Weyand, Stem-like CD4(+) T cells in perivascular tertiary lymphoid structures sustain autoimmune vasculitis. *Sci Transl Med* **15**, eadh0380 (2023).
21. M. Ainciburu, D. M. Morgan, E. A. K. DePasquale, J. C. Love, F. Prosper, P. van Galen, WAT3R: recovery of T-cell receptor variable regions from 3' single-cell RNA-sequencing. *Bioinformatics* **38**, 3645-3647 (2022).

22. I. Sturmlechner, A. Jain, B. Hu, R. R. Jadhav, W. Cao, H. Okuyama, L. Tian, C. M. Weyand, J. J. Goronzy, Aging trajectories of memory CD8 (+) T cells differ by their antigen specificity. *bioRxiv*, (2024).
23. A. Schnell, L. Huang, M. Singer, A. Singaraju, R. M. Barilla, B. M. L. Regan, A. Bollhagen, P. I. Thakore, D. Dionne, T. M. Delorey, M. Pawlak, G. Meyer Zu Horste, O. Rozenblatt-Rosen, R. A. Irizarry, A. Regev, V. K. Kuchroo, Stem-like intestinal Th17 cells give rise to pathogenic effector T cells during autoimmunity. *Cell* **184**, 6281-6298 e6223 (2021).

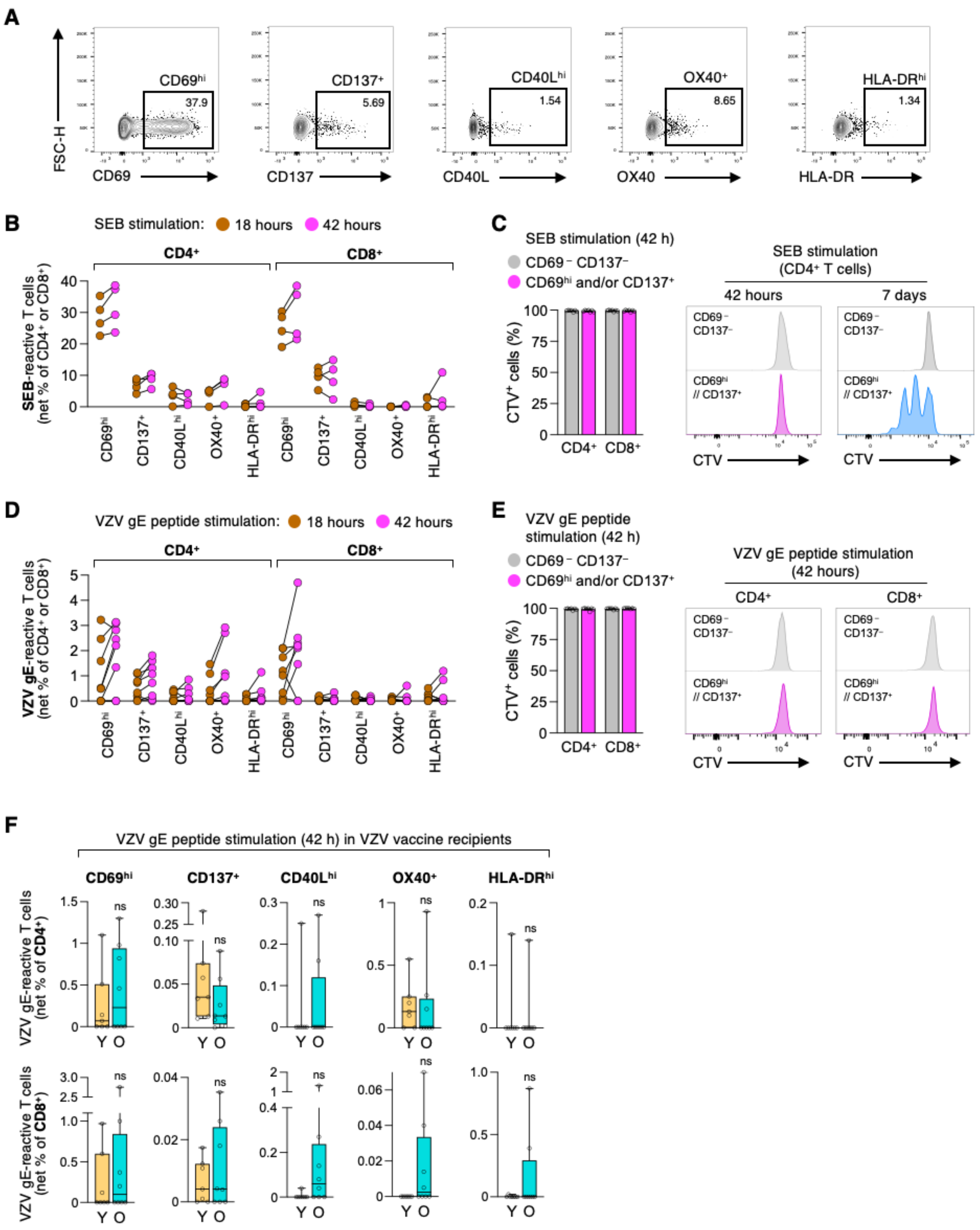
A
FSC-H
CD69hi
37.9
CD137+
5.69
CD40Lhi
1.54
OX40+
8.65
HLA-DRhi
1.34
CD69
CD137
CD40L
OX40
HLA-DR
B
SEB stimulation: 18 hours 42 hours
CD4+
CD8+
SEB-reactive T cells (net % of CD4+ or CD8+)
CD69hi
CD137+
CD40Lhi
OX40+
HLA-DRhi
C
SEB stimulation (42 h)
CD69− CD137−
CD69hi and/or CD137+
CTV+ cells (%)
CD4+
CD8+
SEB stimulation (CD4+ T cells)
42 hours
7 days
CD69− CD137−
CD69hi // CD137+
CTV
D
VZV gE peptide stimulation: 18 hours 42 hours
CD4+
CD8+
VZV gE-reactive T cells (net % of CD4+ or CD8+)
CD69hi
CD137+
CD40Lhi
OX40+
HLA-DRhi
E
VZV gE peptide stimulation (42 h)
CD69− CD137−
CD69hi and/or CD137+
CTV+ cells (%)
CD4+
CD8+
VZV gE peptide stimulation (42 hours)
CD4+
CD8+
CD69− CD137−
CD69hi // CD137+
CTV
F
VZV gE peptide stimulation (42 h) in VZV vaccine recipients
CD69hi
CD137+
CD40Lhi
OX40+
HLA-DRhi
VZV gE-reactive T cells (net % of CD4+)
VZV gE-reactive T cells (net % of CD8+)
ns
Y O

**Supplementary Figure 1, related to Figure 1: Young and older vaccine recipients have similar frequencies of VZV gE-reactive T cells.** The activation-induced marker (AIM) assay was optimized with the goal to comprehensively identify the maximum number of VZV gE-responsive T cells in human peripheral blood. **A**, Representative flow cytometry gating of activation markers after SEB (Staphylococcal Enterotoxin B) stimulation for 42 hours. **B**, SEB-reactive $CD4^+$ and $CD8^+$ T cells in human peripheral blood identified by indicated activation markers at 18 or 42 hours after stimulation. Data show background control-subtracted (net) frequencies. **C,** CellTrace dilution assays to confirm the absence of T cell division at 42 hours after SEB stimulation. A 7-day timepoint was added as positive control. **D**, AIM assay of VZV gE-reactive $CD4^+$ and $CD8^+$ T cells. **E**, CellTrace dilution assays at 42 hours after VZV gE stimulation. **F**, Frequencies of VZV gE-reactive $CD4^+$ and $CD8^+$ T cells in Y and O VZV vaccine recipients as identified by AIM assays. Data show background control-subtracted (net) frequencies. Data show mean ± SEM (C,E) or median (F). All datapoints represent distinct biological replicates. Data were compared by Mann-Whitney tests (F). ns, not significant.

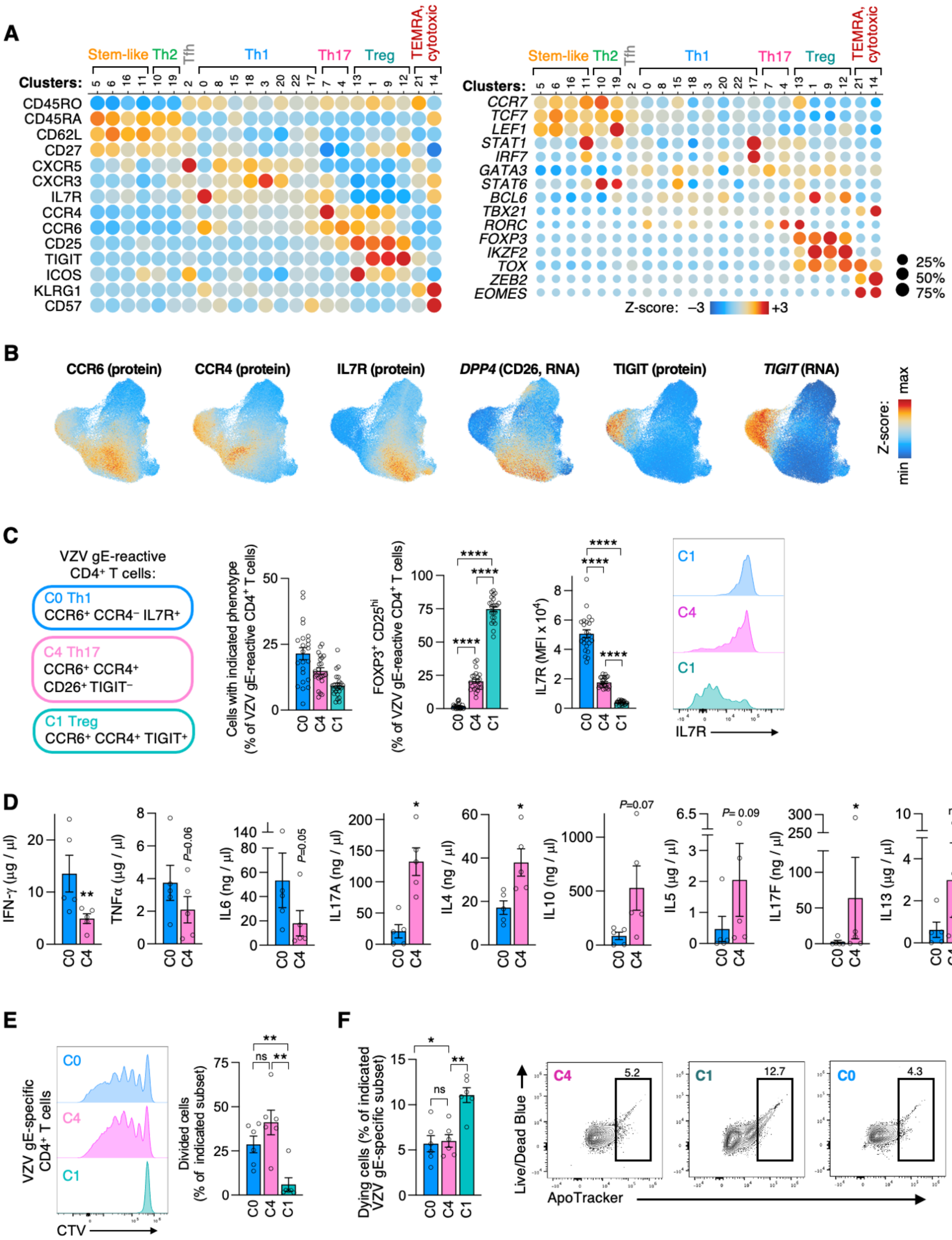
A
Stem-like
Th2
Tfh
Th1
Th17
Treg
TEMRA, cytotoxic
Clusters:
CD45RO
CD45RA
CD62L
CD27
CXCR5
CXCR3
IL7R
CCR4
CCR6
CD25
TIGIT
ICOS
KLRG1
CD57
CCR7
TCF7
LEF1
STAT1
IRF7
GATA3
STAT6
BCL6
TBX21
RORC
FOXP3
IKZF2
TOX
ZEB2
EOMES
Z-score: −3 +3
25%
50%
75%
B
CCR6 (protein)
CCR4 (protein)
IL7R (protein)
DPP4 (CD26, RNA)
TIGIT (protein)
TIGIT (RNA)
Z-score: min max
C
VZV gE-reactive CD4+ T cells:
C0 Th1
CCR6+ CCR4− IL7R+
C4 Th17
CCR6+ CCR4+ CD26+ TIGIT−
C1 Treg
CCR6+ CCR4+ TIGIT+
Cells with indicated phenotype (% of VZV gE-reactive CD4+ T cells)
FOXP3+ CD25hi (% of VZV gE-reactive CD4+ T cells)
IL7R (MFI x 10^4)
IL7R
D
IFN-γ (µg / µl)
TNFα (µg / µl)
IL6 (ng / µl)
IL17A (ng / µl)
IL4 (ng / µl)
IL10 (ng / µl)
IL5 (µg / µl)
IL17F (ng / µl)
IL13 (µg / µl)
P=0.06
P=0.05
P=0.07
P= 0.09
ns
E
VZV gE-specific CD4+ T cells
CTV
Divided cells (% of indicated subset)
F
Dying cells (% of indicated VZV gE-specific subset)
Live/Dead Blue
ApoTracker
5.2
12.7
4.3

**Supplementary Figure 2, related to Figure 1: $CD4^+$ memory T cells specific for VZV gE are phenotypically highly diverse. A,** Dotplot heatmap of protein (left) and gene (right) expression across VZV gE-reactive $CD4^+$ T cell clusters showing classical T cell subset markers. **B**, Feature plots of cluster markers used for FACS experiments. **C**, Flow cytometry of VZV gE-stimulated T cells, examining the expression of FOXP3, CD25, and IL7R in subsets corresponding to clusters C0, C1, C4. **D**, PBMCs of older adults were stimulated with VZV gE peptides. Activated $CD4^+$ T cells with a Th1 (corresponding to C0), Th17 (corresponding to C4), and Treg phenotype (corresponding to C1) were collected. Sorted cells were re-stimulated with αCD3/αCD28 antibodies for 5 days. Cytokine production was measured by LegendPlex multiplex cytokine assays. **E**, VZV gE-reactive $CD4^+$ T cell subsets were collected and stained with CellTrace Violet (CTV) before a 5-day αCD3/αCD28 antibody stimulation. **G**, Proportion of dying cells marked by ApoTracker and viability dye Live/Dead Blue in cultures as in (D). Data show mean ± SEM (C-F). All datapoints represent distinct biological replicates. Data were compared by one-way ANOVA with Tukey's multiple comparisons (C,E,F), two-tailed, paired *t*-test (D). **P*<0.05, ***P*<0.01, ****P*<0.001, *****P*<0.0001. ns, not significant.

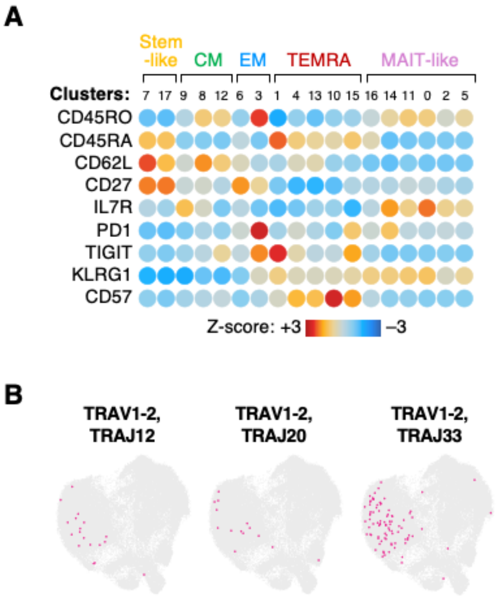


**Supplementary Figure 3, related to Figure 1: The VZV gE-reactive CD8$^{+}$ memory T cell response includes diverse subsets. A,** Dotplot heatmap of CITE protein levels across VZV gE-reactive CD8$^{+}$ T cell clusters showing classical T cell subset markers. **B,** CD8$^{+}$ T cells expressing indicated TRAV-TRAJ segments (pink colored) typically found in MAIT cells.

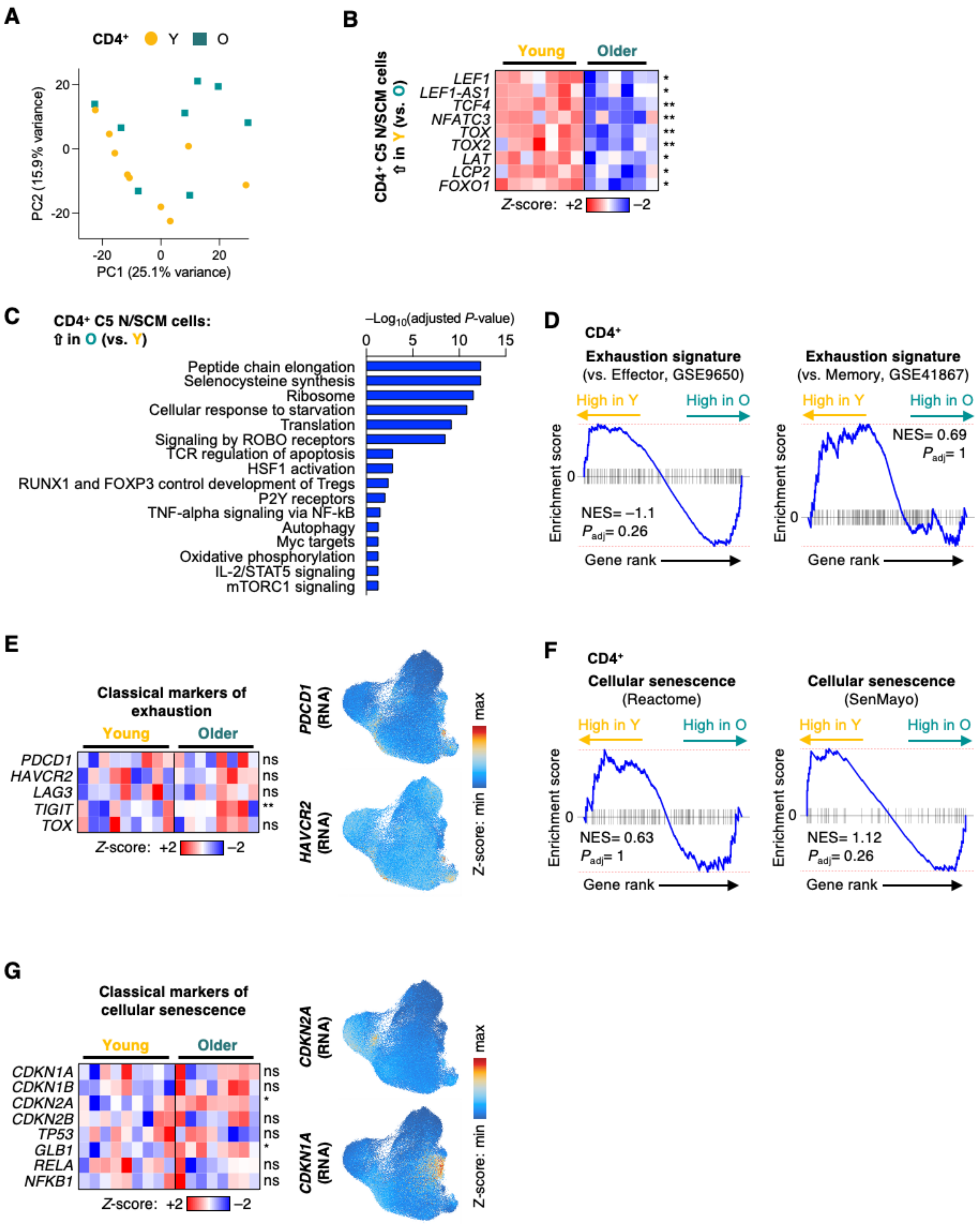


**Supplementary Figure 4, related to Figure 3: Stem-like features in the CD4⁺ T cell memory response from older adults against VZV gE are diminished. A**, Principal component analysis of CD4⁺ VZV gE-specific single cell transcripts aggregated into pseudo-bulk data for each vaccine recipient. **B**, Heatmap of pseudo-bulk gene expression of selected $CD4^+$ T cell DEGs in C5. Only clusters with minimum sample representation of 2 male and 2 female participants per group are included. **C**, Pathway enrichment of $CD4^+$ C5 DEGs higher expressed in O than in Y. **D**, GSEA of gene expression in O vs Y for T cell exhaustion gene sets. **E**, Heatmap of pseudo-bulk gene expression (left) and feature plots (right) of classical markers of T cell exhaustion. **F**, GSEA for cellular senescence gene sets. **G**, Heatmap and feature plots for classical cellular senescence markers.

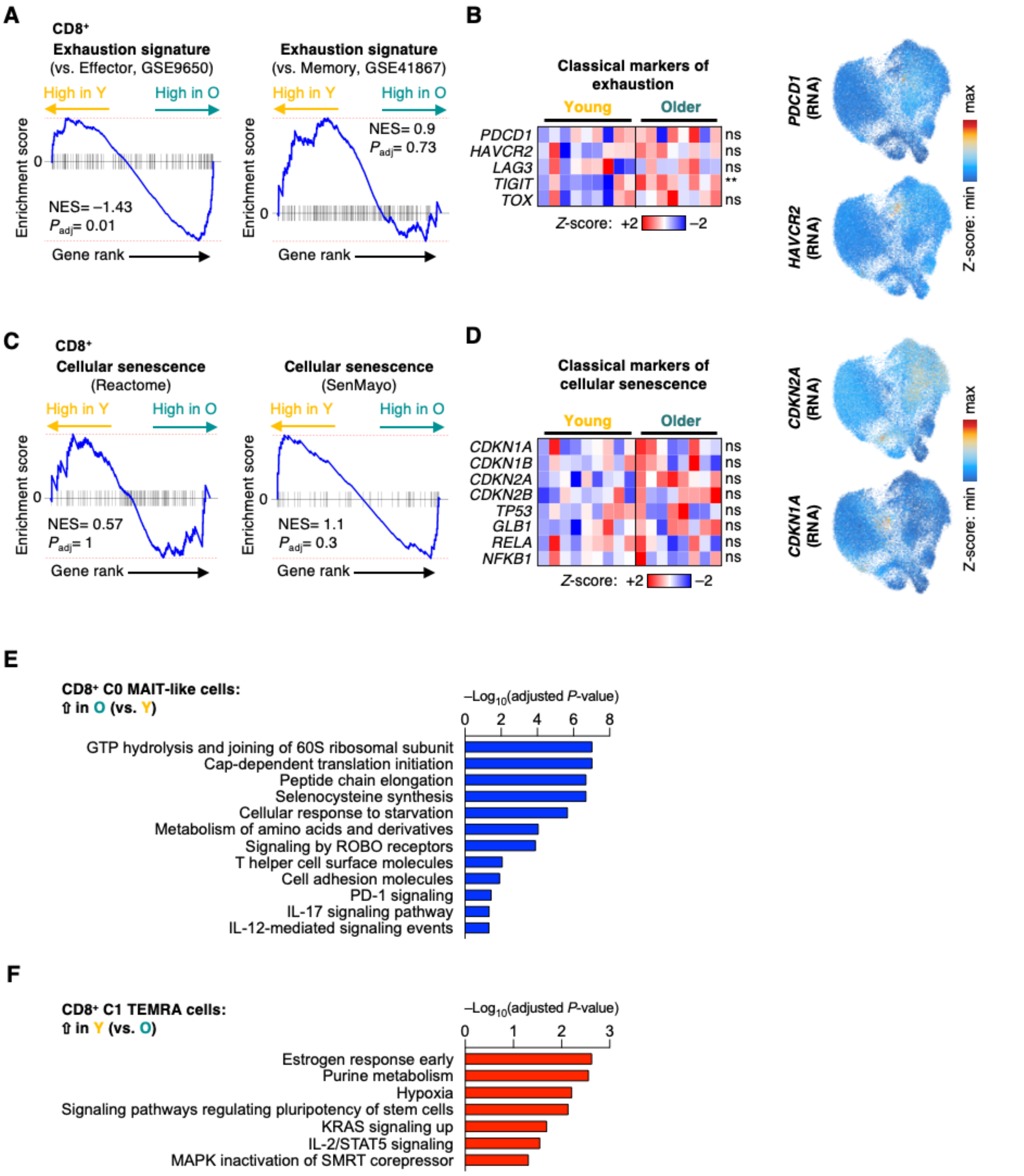


**Supplementary Figure 5, related to Figure 3: CD8⁺ memory T cells specific to VZV gE lack signatures of age-related exhaustion or cellular senescence. A**, GSEA of T cell exhaustion gene sets for pseudo-bulk expression of total VZV gE-reactive CD8⁺ T cells in Y versus O vaccine recipients. **B**, Heatmap of pseudo-bulk gene expression (left) and feature plots (right) of classical markers of T cell exhaustion. **C**, GSEA of cellular senescence gene sets as in (A). **D**, Heatmap of pseudo-bulk gene expression (left) and feature plots (right) of classical cellular senescence markers. **E**, Pathway enrichment for $CD8^+$ C0 DEGs higher expressed in O than in Y. **F**, Pathway enrichment for $CD8^+$ C1 DEGs higher expressed in Y than in O.

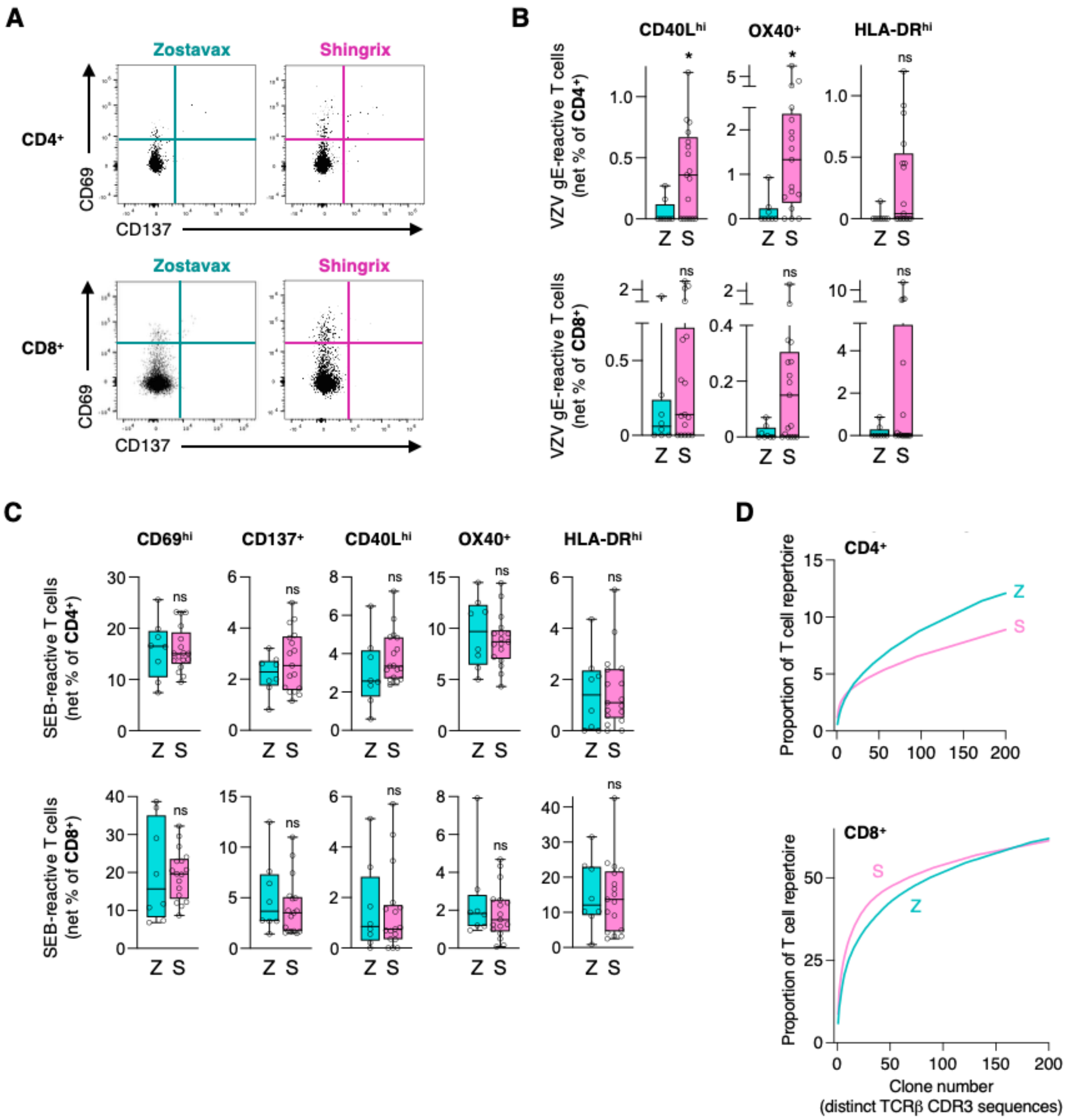


**Supplementary Figure 6, related to Figure 4: VZV gE responses are increased in Shingrix recipients. A**, Representative flow cytometry plot of CD69$^{hi}$ and/or CD137$^{+}$ T cells after VZV gE peptide stimulation. **B**, Proportion of VZV gE-reactive CD4$^{+}$ or CD8$^{+}$ T cells in Z or S vaccine recipients as identified by flow cytometry for activation markers CD40L$^{hi}$, OX40$^{hi}$ or HLA-DR$^{hi}$. Data show background control-subtracted (net) frequencies. **C**, Proportion of SEB-reactive CD4$^{+}$ or CD8$^{+}$ T cells in Z or S vaccine recipients as identified by indicated activation marker in flow cytometry analysis. Data show background control-subtracted (net) frequencies. **D**, Cumulative

TCR frequency plots for VZV gE-specific $CD4^+$ (top) and $CD8^+$ T cells (bottom) from Z and S recipients in single cell sequencing experiments. The plot shows TCRs ranked by t descending clone sizes versus the space they occupy. Data show median (B,D). All datapoints represent distinct biological replicates. Data were compared by Mann-Whitney tests (B,C). *$P$<0.05. ns, not significant.

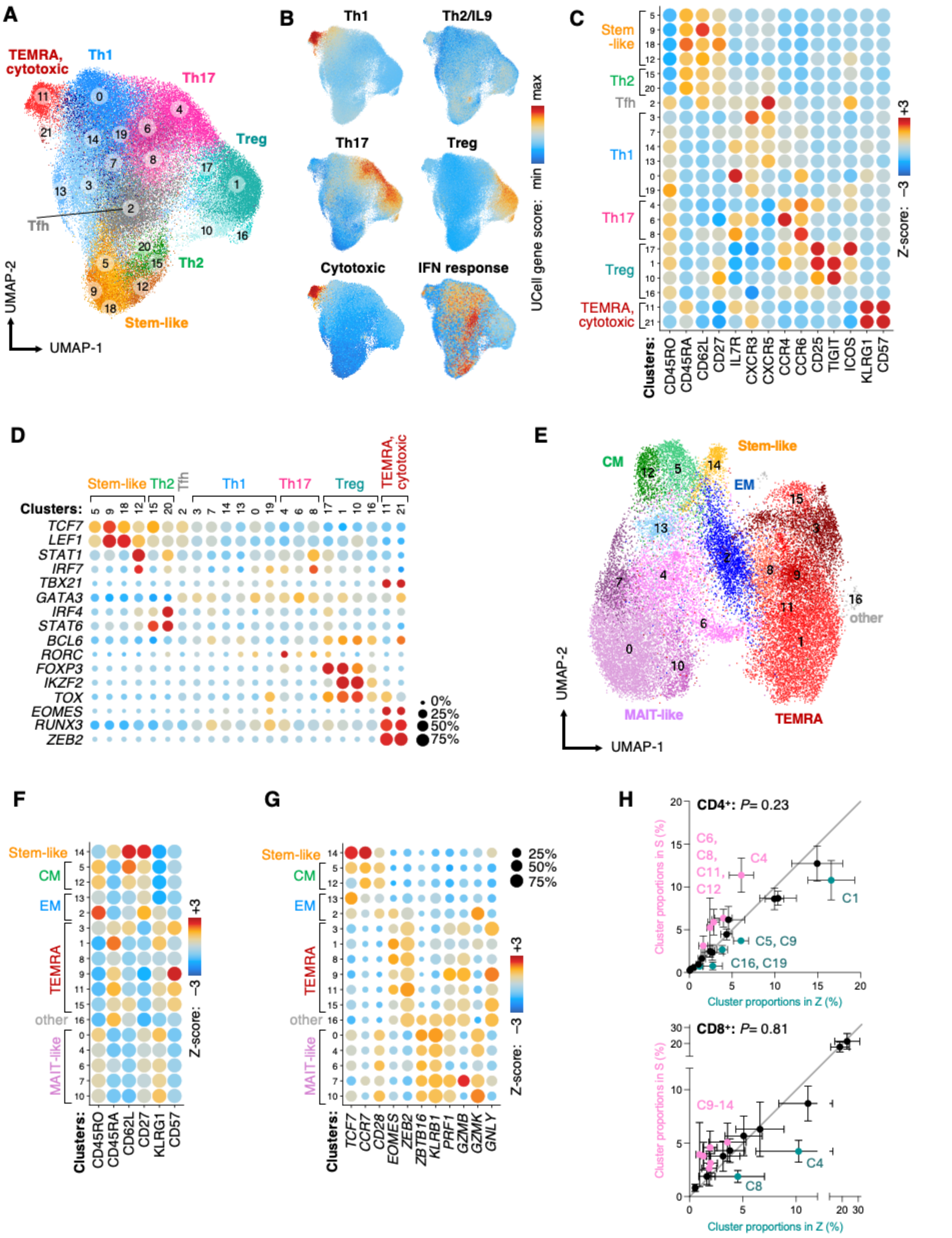
A
TEMRA, cytotoxic
Th1
Th17
Treg
Tfh
Th2
Stem-like
UMAP-2
UMAP-1
B
Th1
Th2/IL9
Th17
Treg
Cytotoxic
IFN response
UCell gene score: min max
C
Stem -like
Th2
Tfh
Th1
Th17
Treg
TEMRA, cytotoxic
Clusters:
CD45RO CD45RA CD62L CD27 IL7R CXCR3 CXCR5 CCR4 CCR6 CD25 TIGIT ICOS KLRG1 CD57
Z-score: −3 +3
D
Stem-like Th2 Tfh Th1 Th17 Treg TEMRA, cytotoxic
Clusters:
TCF7 LEF1 STAT1 IRF7 TBX21 GATA3 IRF4 STAT6 BCL6 RORC FOXP3 IKZF2 TOX EOMES RUNX3 ZEB2
0% 25% 50% 75%
E
CM
Stem-like
EM
other
MAIT-like
TEMRA
UMAP-2
UMAP-1
F
Stem-like
CM
EM
TEMRA
other
MAIT-like
Clusters:
CD45RO CD45RA CD62L CD27 KLRG1 CD57
Z-score: −3 +3
G
Stem-like
CM
EM
TEMRA
other
MAIT-like
Clusters:
TCF7 CCR7 CD28 EOMES ZEB2 ZBTB16 KLRB1 PRF1 GZMB GZMK GNLY
25% 50% 75%
Z-score: −3 +3
H
CD4⁺: P= 0.23
C6, C8, C11, C12
C4
C1
C5, C9
C16, C19
Cluster proportions in S (%)
Cluster proportions in Z (%)
CD8⁺: P= 0.81
C9-14
C4
C8
Cluster proportions in S (%)
Cluster proportions in Z (%)

**Supplementary Figure 7, related to Figure 4: VZV gE-specific T cells in Shingrix and Zostavax vaccine recipients include diverse subsets. A**, UMAP of VZV gE-reactive CD4$^+$ T cells of S and Z vaccine recipients. **B**, Feature plots of CD4$^+$ T cell UCell gene scores supporting CD4$^+$ T cell subset annotation. **C**, Bubble plot of CITE protein expression of classical T cell subset markers across CD4$^+$ T cell clusters. **D**, Bubble plots of gene expression across CD4$^+$ T cell clusters showing subset-specific transcription factors (left) and cytokines (right). The bubble size corresponds to the proportion of cells that express a given gene. **E**, UMAP of VZV gE-reactive CD8$^+$ T cells of S and Z vaccine recipients. **F**, Bubble plot of CITE protein expression across CD8$^+$ T cell clusters showing classical T cell subset markers. **G**, Bubble plot of gene expression across CD8$^+$ T cell clusters showing classical T cell subset markers. The bubble size corresponds to the proportion of cells that express a given gene. **H**, Cluster distribution of CD4$^+$ or CD8$^+$ VZV gE-reactive T cells from Z and S vaccine recipients. Statistical analysis was performed comparing probability vectors and permutation tests.

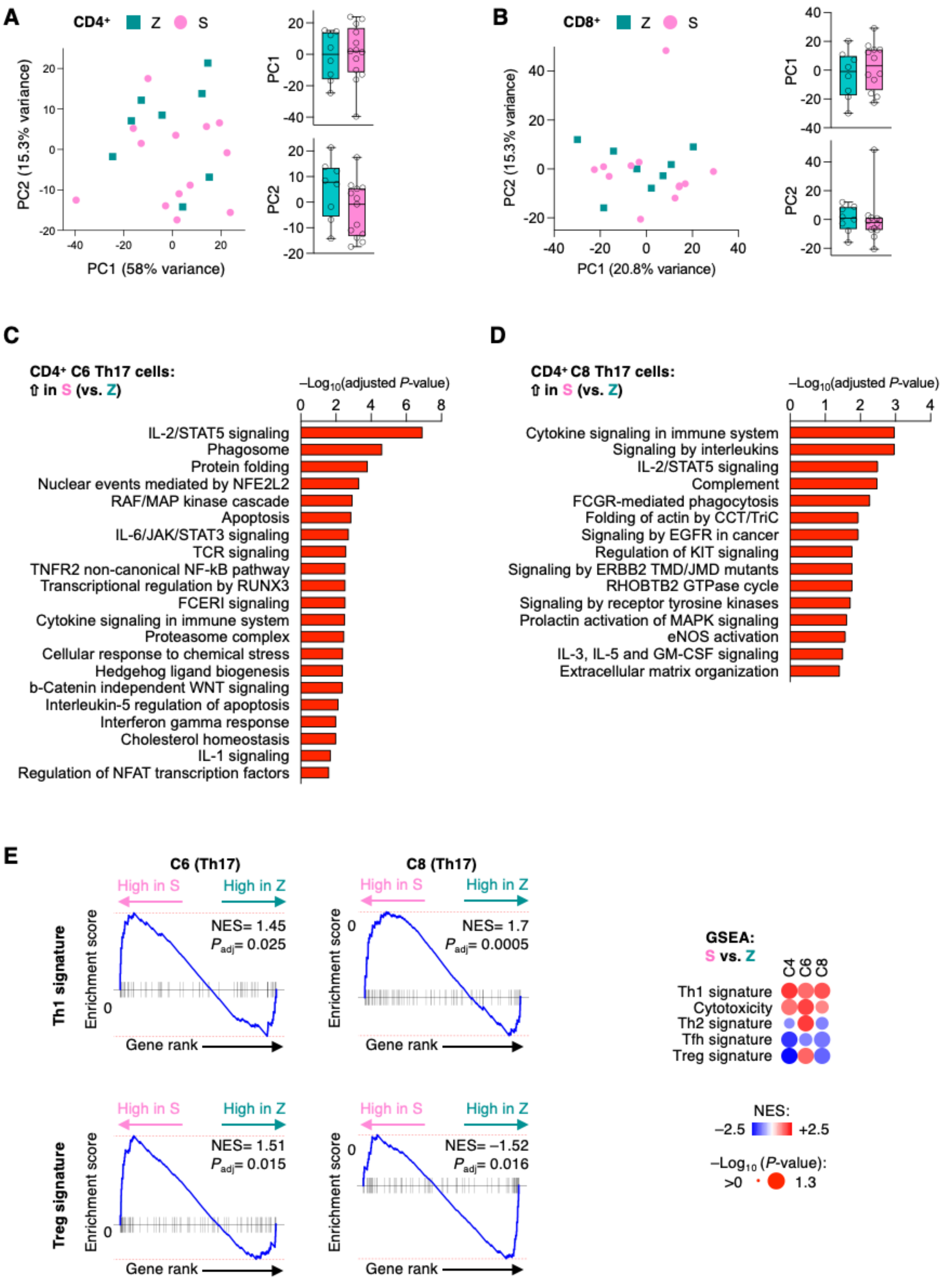
A
CD4+ Z S
PC2 (15.3% variance)
PC1 (58% variance)
PC1
PC2
B
CD8+ Z S
PC2 (15.3% variance)
PC1 (20.8% variance)
PC1
PC2
C
CD4+ C6 Th17 cells:
⇧ in S (vs. Z)
–Log10(adjusted P-value)
IL-2/STAT5 signaling
Phagosome
Protein folding
Nuclear events mediated by NFE2L2
RAF/MAP kinase cascade
Apoptosis
IL-6/JAK/STAT3 signaling
TCR signaling
TNFR2 non-canonical NF-kB pathway
Transcriptional regulation by RUNX3
FCERI signaling
Cytokine signaling in immune system
Proteasome complex
Cellular response to chemical stress
Hedgehog ligand biogenesis
b-Catenin independent WNT signaling
Interleukin-5 regulation of apoptosis
Interferon gamma response
Cholesterol homeostasis
IL-1 signaling
Regulation of NFAT transcription factors
D
CD4+ C8 Th17 cells:
⇧ in S (vs. Z)
–Log10(adjusted P-value)
Cytokine signaling in immune system
Signaling by interleukins
IL-2/STAT5 signaling
Complement
FCGR-mediated phagocytosis
Folding of actin by CCT/TriC
Signaling by EGFR in cancer
Regulation of KIT signaling
Signaling by ERBB2 TMD/JMD mutants
RHOBTB2 GTPase cycle
Signaling by receptor tyrosine kinases
Prolactin activation of MAPK signaling
eNOS activation
IL-3, IL-5 and GM-CSF signaling
Extracellular matrix organization
E
C6 (Th17)
C8 (Th17)
High in S
High in Z
Th1 signature
Enrichment score
NES= 1.45
Padj= 0.025
NES= 1.7
Padj= 0.0005
Gene rank
Treg signature
NES= 1.51
Padj= 0.015
NES= –1.52
Padj= 0.016
GSEA:
S vs. Z
C4 C6 C8
Th1 signature
Cytotoxicity
Th2 signature
Tfh signature
Treg signature
NES:
–2.5 +2.5
–Log10 (P-value):
>0 1.3

**Supplementary Figure 8, related to Figure 6: VZV gE-reactive CD4$^+$ T cells in all Th17-related clusters share gene expression profiles indicating increased functionality in Shingrix recipients. A-B**, Principal component analysis on pseudo-bulk transcripts of VZV gE-reactive CD4$^+$ (A) and CD8$^+$ (B) T cells from Z and S vaccine recipients (left). PC-1 and 2 are shown as box plots with medians. **C-D**, Pathway enrichment for CD4$^+$ clusters C6 (C) and C8 (D) of DEGs higher expressed in S than in Z. **E**, GSEA of C6 and C8 pseudo-bulk gene expression in Z versus S for concordance with Th1 and Treg gene sets. NES, normalized enrichment score.

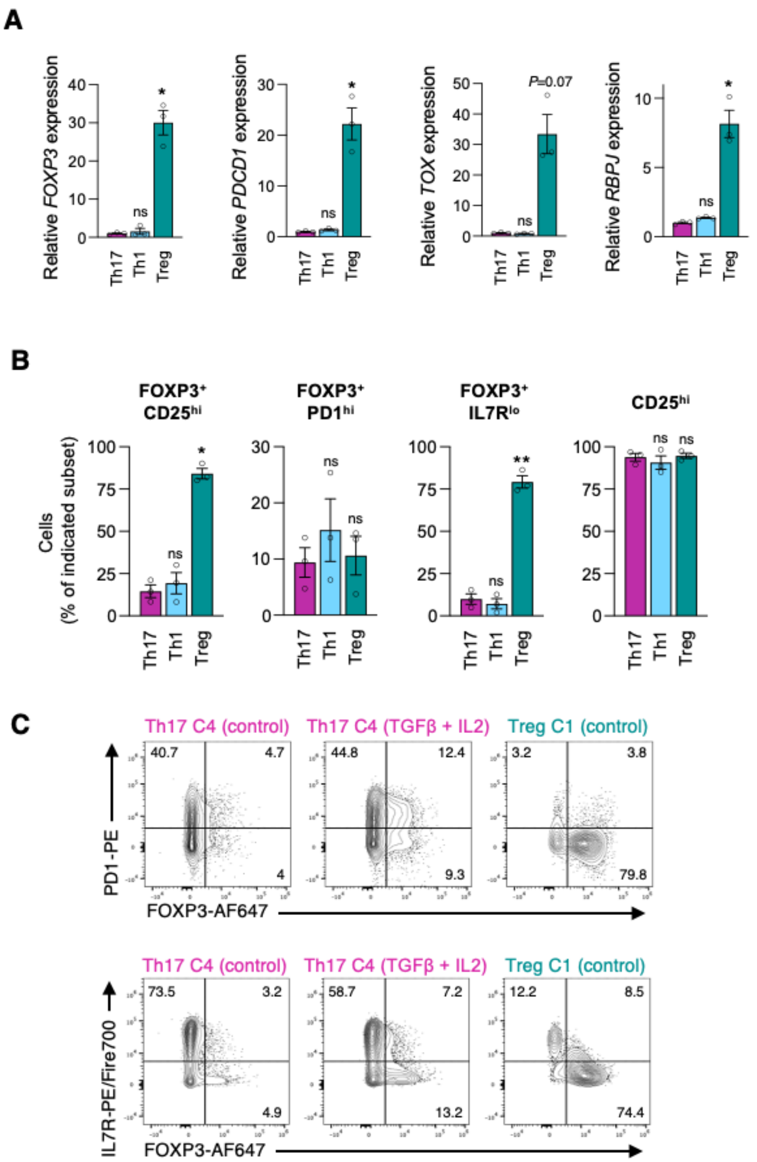


**Supplementary Figure 9, related to Figure 6: Signatures of Th17 cells overlap with cluster 1 Treg phenotypes under selective conditions. A**, qPCR quantitative analysis of cells with a cluster 4 Th17, cluster 0 Th1 or cluster 1 Treg phenotype that were FACS-collected from peripheral memory $CD4^+$ T cells of older adults and stimulated with αCD3/αCD28 antibodies for 3 days. Data are representative of 3 experiments. **B**, Flow cytometry quantitation of cells as in (A) but stimulated for 7 days. **C**, Flow cytometry plots of Th17 or Treg cells as in (A) that were stimulated with αCD3/αCD28 in the absence of presence of 25 ng/ml TGFβ and 500 U/ml IL2 for

7 days. Data show mean ± SEM (A,B). All datapoints represent distinct biological replicates. Data were compared by one-way ANOVA with Šídák's multiple comparisons test (A,B). *$P<0.05$, **$P<0.01$. ns, not significant.

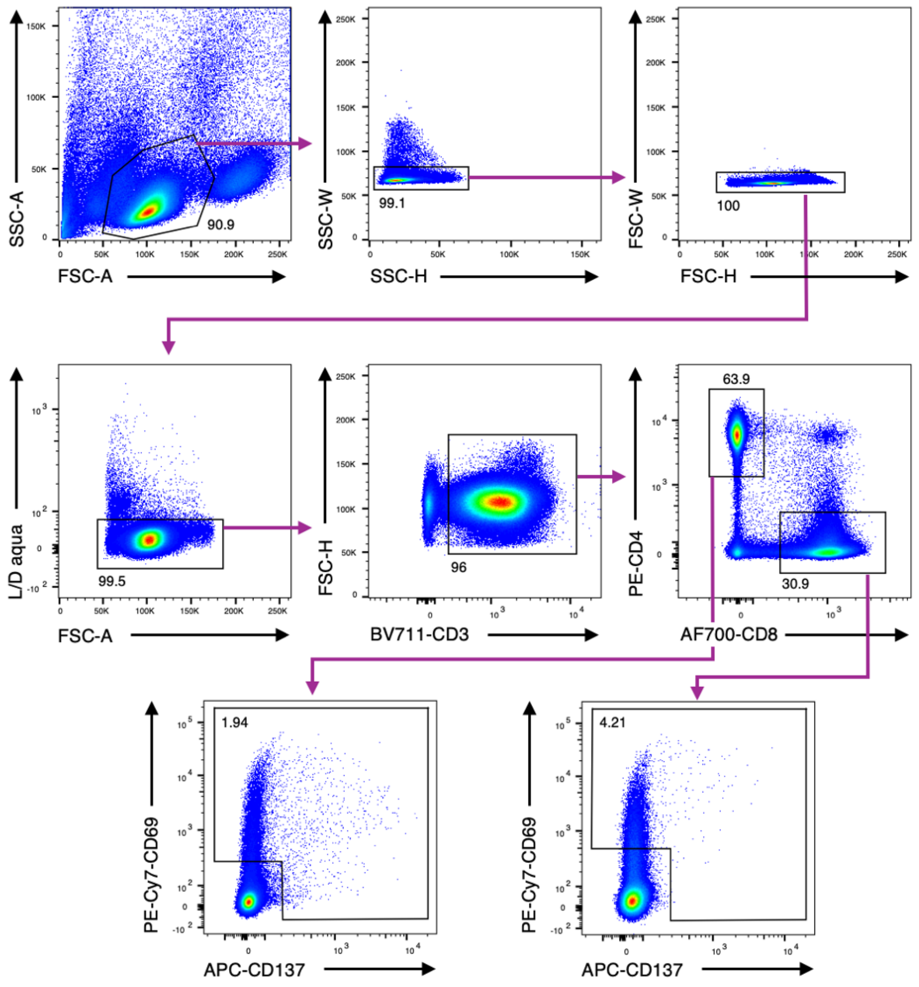


**Flow Cytometry Gating Strategy 1: FACS gating strategy to collect VZV gE-reactive T cells for single cell sequencing.**

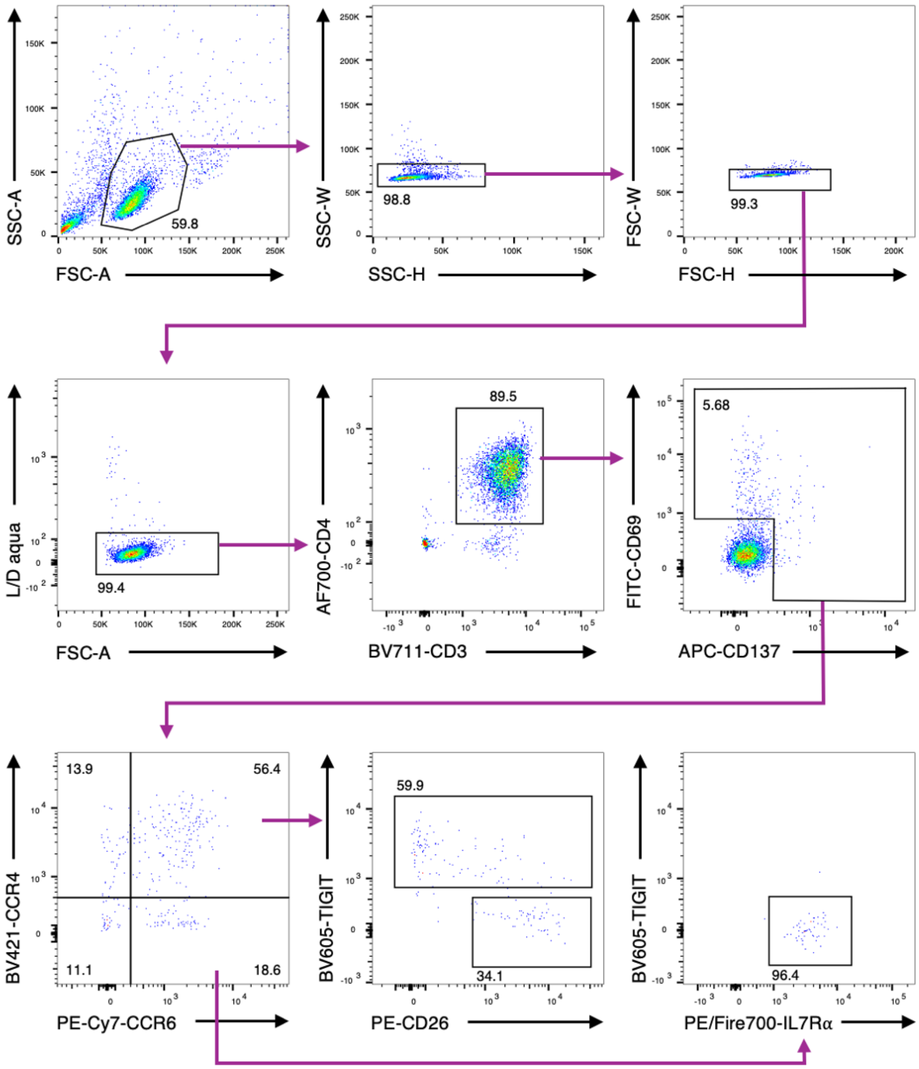


**Flow Cytometry Gating Strategy 2: FACS gating strategy to collect phenotypic subsets of VZV gE-reactive T cells for functional studies.**

**Supplementary Table 1: Patient demographics for single cell sequencing samples**

| | |
|---|---|
| Volunteers<br>for single cell sequencing | 30 |
| Female | 46.7% |
| Male | 53.3% |
| White | 83.3% |
| Asian | 13.3% |
| American Indian/Alaska Native | 3.3% |
| Race Unknown / Not Reported | 3.3% |
| Not Hispanic or Latino | 96.6% |
| Ethnicity Unknown / Not Reported | 3.3% |

| | Varivax, Young | Zostavax, Older | Shingrix, Older |
|---|---|---|---|
| Volunteers | 9 | 8 | 13 |
| Age at sample collection (years) | 24.3 ± 2.9 | 75.5 ± 7.3 | 69.5 ± 7.3 |
| Age at last VZV vaccination (years) | 9.8 ± 3.3 | 69.5 ± 7.6 | 65.6 ± 7.4 |
| Years since last vaccination (years) | 14.4 ± 1.1 | 6 ± 0.8 | 3.4 ± 0.5 |

**Supplementary Table 2: Patient comorbidities and key medication for single cell sequencing samples**

| Volunteers for single cell sequencing | Zostavax | Shingrix |
|---|---|---|
| Total number | 8 | 13 |
| Gender (M/F) | 4/4 | 8/5 |
| Major comorbidities | 3 (1-4) | 3 (1-5) |
| Diabetes mellitus | 3 (38%) | 3 (23%) |
| Atrial fibrillation | 2 (25%) | 6 (46%) |
| Chronic kidney disease | 1 (13%) | 3 (23%) |
| Coronary artery disease | 1 (13%) | 2 (15%) |
| Hypertension | 3 (38%) | 5 (38%) |
| Hyperlipidemia | 5 (63%) | 9 (69%) |
| Asthma | 1 (13%) | 3 (23%) |
| COPD | 2 (25%) | 0 (0%) |
| Medication | | |
| Statin | 4 (50%) | 8 (62%) |
| Anti-diabetics | 2 (25%) | 3 (23%) |
